# Title: β Cell microRNAs Function as Molecular Hubs of Type 1 Diabetes Pathogenesis and as Biomarkers of Diabetes Risk

**DOI:** 10.1101/2023.06.15.545170

**Authors:** Farooq Syed, Preethi Krishnan, Garrick Chang, Sarah R. Langlais, Sumon Hati, Kentaro Yamada, Anh K. Lam, Sayali Talware, Xiaowen Liu, Rajesh Sardar, Jing Liu, Raghavendra G. Mirmira, Carmella Evans-Molina

## Abstract

MicroRNAs (miRNAs) are small non-coding RNAs that play a crucial role in modulating gene expression and are enriched in cell-derived extracellular vesicles (EVs). We investigated whether miRNAs from human islets and islet-derived EVs could provide insight into β cell stress pathways activated during type 1 diabetes (T1D) evolution, therefore serving as potential disease biomarkers. We treated human islets from 10 cadaveric donors with IL-1β and IFN-γ to model T1D *ex vivo*. MicroRNAs were isolated from islets and islet-derived EVs, and small RNA sequencing was performed. We found 20 and 14 differentially expressed (DE) miRNAs in cytokine-versus control-treated islets and EVs, respectively. Interestingly, the miRNAs found in EVs were mostly different from those found in islets. Only two miRNAs, miR-155-5p and miR-146a-5p, were upregulated in both islets and EVs, suggesting selective sorting of miRNAs into EVs. We used machine learning algorithms to rank DE EV-associated miRNAs, and developed custom label-free Localized Surface Plasmon Resonance-based biosensors to measure top ranked EVs in human plasma. Results from this analysis revealed that miR-155, miR-146, miR-30c, and miR-802 were upregulated and miR-124-3p was downregulated in plasma-derived EVs from children with recent-onset T1D. In addition, miR-146 and miR-30c were upregulated in plasma-derived EVs of autoantibody positive (AAb+) children compared to matched non-diabetic controls, while miR-124 was downregulated in both T1D and AAb+ groups. Furthermore, single-molecule fluorescence in situ hybridization confirmed increased expression of the most highly upregulated islet miRNA, miR-155, in pancreatic sections from organ donors with AAb+ and T1D.

**One Sentence Summary:** miRNA expression patterns in human pancreatic islets and EVs change under inflammatory conditions and can be leveraged to inform biomarkers strategies for T1D.

## INTRODUCTION

Type 1 diabetes (T1D) results from immune-mediated destruction of pancreatic β cells [1]. Data from natural history studies enrolling newborns with high genetic risk indicate that nearly 70% of individuals with two or more T1D-associated autoantibodies will develop clinical disease within 10 years of follow-up [2]. These epidemiological data formed the basis for the delineation of a new staging paradigm in 2015, where Stage 1 T1D is defined as the presence of two or more islet autoantibodies and normal glucose tolerance, Stage 2 disease is defined as the presence of multiple islet autoantibodies and dysglycemia, and Stage 3 T1D is marked by the development of hyperglycemia that exceeds the American Diabetes Association thresholds for clinical diagnosis [3–7]. Immunomodulatory interventions initiated at the onset of Stage 3 T1D have shown limited efficacy in inducing a durable disease remission [4–8]. However, teplizumab, an Fc receptor– nonbinding anti-CD3 monoclonal antibody, delayed the onset of Stage 3 T1D by approximately 32 months in high-risk individuals in Stage 2 T1D [9,10]. Teplizumab was approved by the United States Food and Drug Administration (FDA) in November 2022 as the first disease-modifying therapy for T1D.

While the teplizumab trial and approval provides the first evidence that immune interventions are able to substantially alter the course of T1D, variable responses to disease-modifying therapies has been observed in all clinical trials performed to date [11]. In addition, not all individuals with autoantibodies will progress to clinical disease, and not all autoantibodies are associated with the same risk of disease progression [12]. Thus, the timeframe of T1D development in at-risk individuals is highly variable, which leads to challenges in timing therapeutic interventions. Biomarkers capable of stratifying heterogeneous at-risk populations, especially markers that provide insight into the health status of the β cell, have the potential to identify individuals most likely to benefit from early immunomodulatory intervention. In addition, biomarkers may also point to molecular pathways associated with disease pathogenesis and therefore help inform new approaches for disease interdiction.

MicroRNAs (miRNAs) are a class of endogenously produced and highly-conserved small non-coding RNAs that are ∼22 nucleotides in length. The most well-known mechanism of action of miRNAs is inhibition of gene expression through complementary binding to the 3’ untranslated region of mRNAs to either repress mRNA translation or cause mRNA degradation. A broad range of biological processes within the β cell are regulated by miRNAs, including insulin secretion, endoplasmic reticulum (ER) stress, and apoptosis [13–16]. Notably, miRNAs can be detected in a variety of body fluids, including serum, plasma, urine, saliva, and breastmilk [17–22]. Recent evidence suggests that miRNAs are enriched in extracellular vesicles (EVs), a class of membrane-bound vesicles that serve as important mediators of intracellular communication [20–22].

In the current study, we hypothesized that miRNA signatures of islet and islet-derived EVs could be leveraged as biomarkers to detect ongoing β cell stress during the evolution of T1D, while also offering insight into β cell pathways linked with disease development. To test this hypothesis, we first utilized an *ex vivo* human islet-based model of inflammatory T1D stress to perform unbiased miRNA discovery experiments. Next, we used custom-made, label-free LSPR-based biosensors to validate differentially expressed (DE) miRNA signatures in plasma EVs isolated from children with autoantibody positivity (AAb+) or T1D or healthy controls. Finally, we verified upregulation of miR-155 in the pancreas of organ donors with AAb+ and T1D using single molecule RNA fluorescence in situ hybridization (FISH).

## RESULTS

### Isolation and characterization of human islet-derived extracellular vesicles

Fig. 1 illustrates our experimental workflow. In brief, (A) human islets were treated with or without IL-1β + IFN-γ as an *ex vivo* model of T1D, followed by small RNA sequencing of human islet and islet-derived EVs; (B) DE miRNAs were validated in separate cytokine-treated islet batches using targeted expression analysis; (C) prioritized EV-miRNAs were measured in plasma-derived EVs from individuals with AAb+ and recent onset T1D and matched healthy controls; (D) miR-155 expression patterns were analyzed in pancreatic samples from human organ donors with AAb+ and T1D and non-diabetic controls.

**Figure 1:**
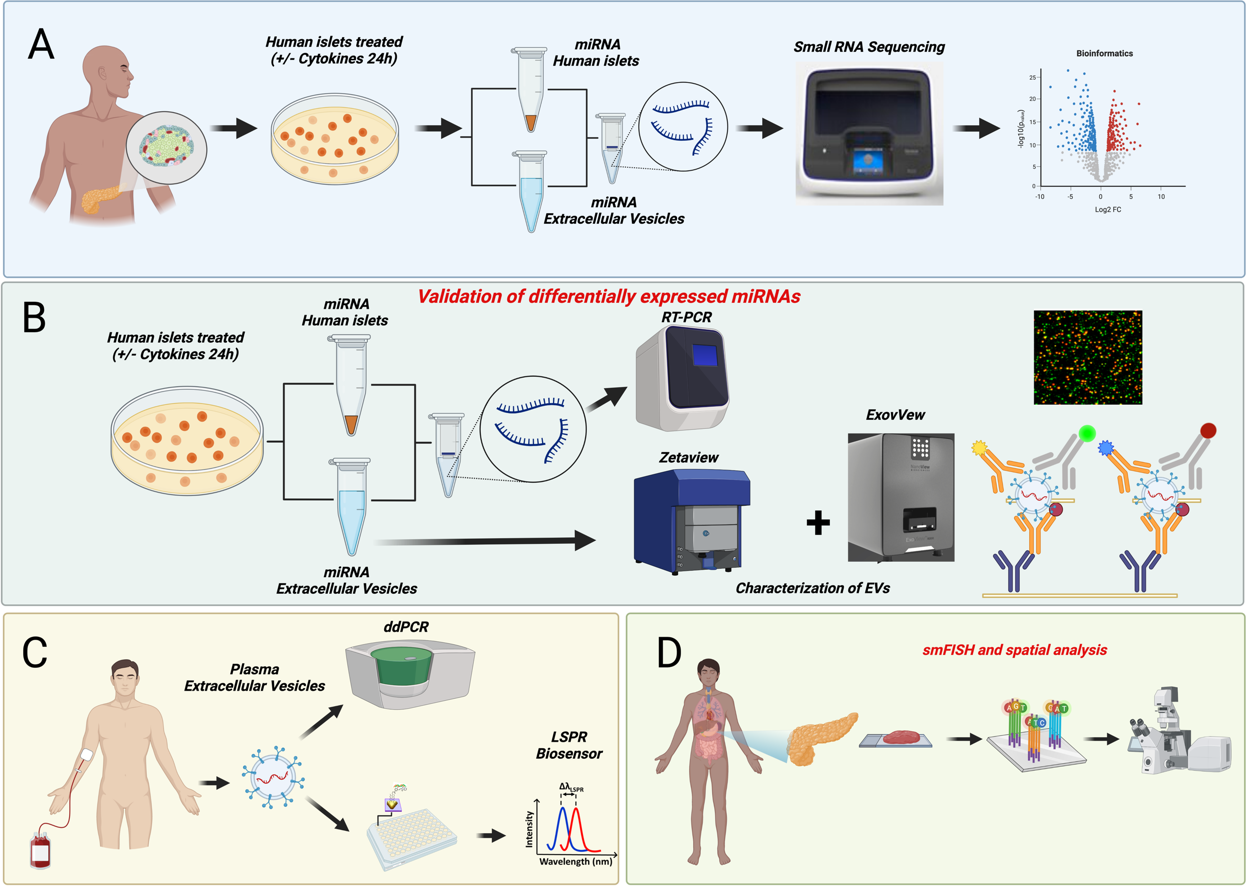
Schematic representation of the experimental workflow. Islets obtained from human cadaveric donors (n=10 total; 5M/5F) were treated with IL-1β+IFN-γ for 24 hrs. (A) Total RNA was isolated from islets and islet-derived EVs and subjected to small RNA sequencing. (B) Differentially expressed miRNAs were further validated using separate sets of islets and islet-derived EVs. (C) A panel of miRNAs selected from the top five differentially expressed EV miRNAs were validated using human circulating plasma-derived EVs from healthy controls, individuals that are AAb+, and individuals with recent onset T1D. (D) miR-155 expression patterns were analyzed in pancreatic samples from human organ donors with AAb+ and T1D and non-diabetic controls.

In human islet cultures treated with or without IL-1β + IFN-γ, EVs were isolated using a proprietary ultracentrifugation-free method, and EVs were assessed using transmission electron microscopy (TEM), scanning electron microscopy (SEM), qRT-PCR, Nanoparticle Tracking Analysis (NTA), immunoblot, and single particle interferometric reflected imaging. Human islets treated with or without cytokines showed similar ultra-structural morphology of intracellular multivesicular bodies (MVBs) as determined by TEM analysis (Fig. 2A), and SEM analysis confirmed the presence of EV particles in the plasma membrane (Fig. 2B). SEM and TEM analysis of EVs isolated from control and cytokine-treated human islet conditioned media confirmed the presence of EVs, and similar ultrastructural features of EVs were noted under both conditions (Fig 2C).

**Figure 2:**
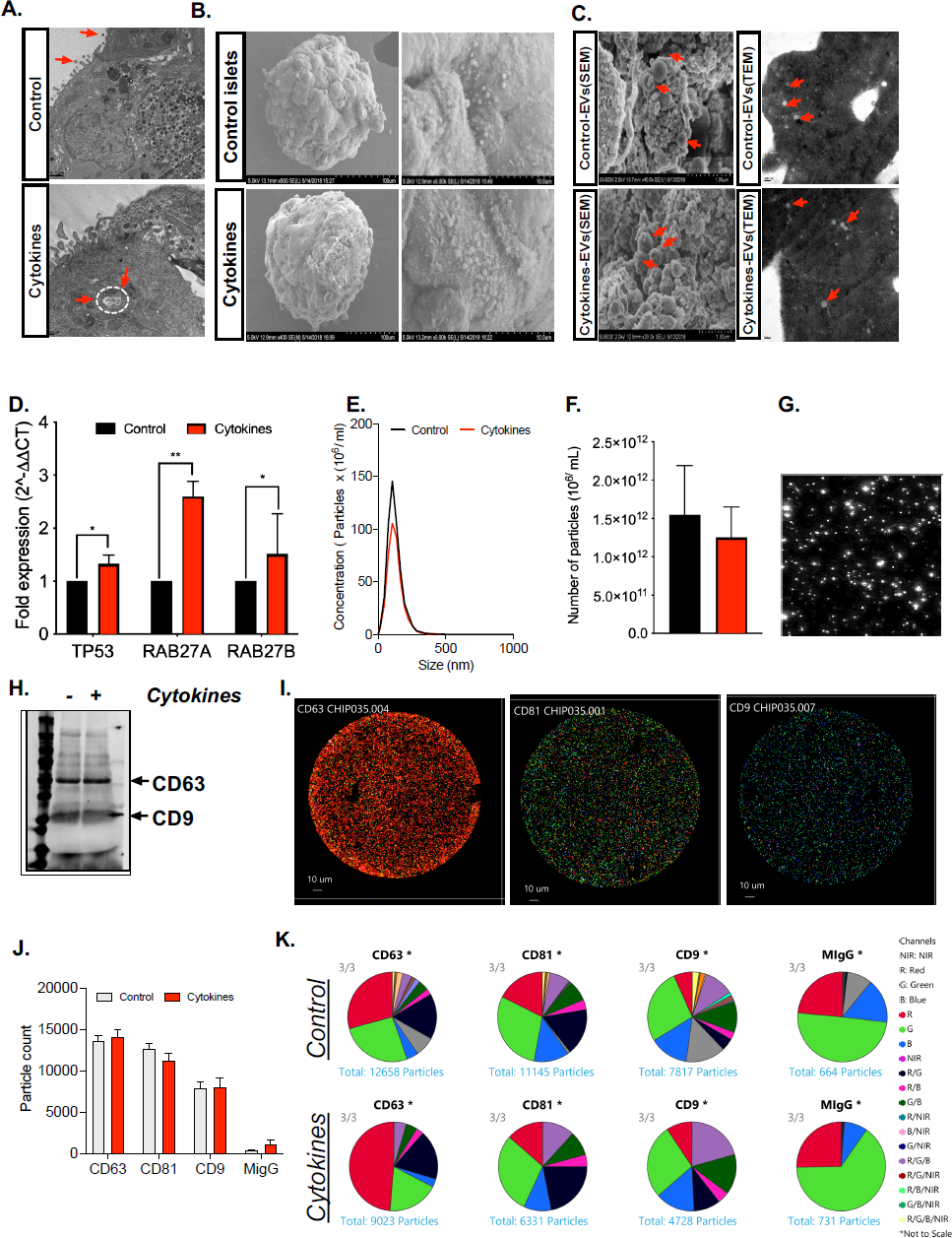
Characterization of isolated EVs. Human islets were treated with or without 50 U/mL IL-1β and 1000 U/mL IFN-γ for 24 hours. (A) Islets were then subjected to TEM analysis to identify the presence of EVs over the islet plasma membrane and multivesicular bodies (MVs) inside the cytoplasmic region of pancreatic β cells, indicated by red arrows. (B) SEM images of human islets show the presence of EVs over the islet plasma membrane (C) Representative SEM and TEM images of isolated EVs from control and cytokine-treated islet culture supernatant. (D) qRT-PCR analysis of markers involved in EV biogenesis in control and cytokine-treated islets (*p<0.05, **p<0.01. n = 6). (E-F) nanoparticle tracking analysis (NTA) showing size distribution graphs for isolated EVs from control (black line) and cytokine-treated (red line) islets. (G) Bright field images from NTA. (H) Immunoblot analysis for CD63 and CD9 from isolated EVs. (I) Images of Exoview showing the presence of EVs captured using CD63, CD81, CD9 and mouse IgG(MIgG) was used as an negative control. (J) Quantification of EV particles and (K) co-localization of tetraspanin marker with each other markers from control and cytokine treated human islet culture supernatant. (n = 3).

Cytokine treatment increased expression of several transcripts involved in EV biogenesis, including TP53, RAB27A, and RAB27B (Fig. 2D). However, NTA showed that EVs exhibited a uniform size range of 50-200 nm with no significant difference in particle concentration or size distribution between EVs secreted from control or cytokine-treated human islets (Fig. 2E-G). Immunoblot analysis of isolated EVs confirmed the presence of the tetraspanin EV markers CD63 and CD9 (Fig. 2H). To further characterize tetraspanin distribution patterns, EVs were isolated by dual size exclusion chromatography and characterized using single particle interferometric reflected imaging (Fig. 2I). Results from this analysis showed that a higher proportion of islet-derived EVs were positive for CD63 compared to CD9 and CD81 (Fig. 2J). Co-localization analysis of tetraspanin markers CD63, CD9, and CD81 using a chip-based EV analysis (Exoview) showed no difference between EVs isolated from control and cytokine-treated islet supernatant (Fig. 2K), indicating that the composition of EV markers did not change dramatically between control and cytokine conditions.

### Inflammatory cytokine treatment leads to differential expression of miRNAs in human islets and islet-derived EVs

Next, RNA was isolated from control and cytokine-treated islets and islet-derived EVs and subjected to unbiased small RNA sequencing. A total of 83,141,746 and 87,979,932 reads were detected in samples obtained from control and cytokine-treated islets, respectively. Of these, ∼65% of the reads in control and 67.5% of the reads in cytokine-treated islets aligned to the reference genome. Approximately one-third of the aligned reads (∼32%) mapped to mature miRNAs in both control and cytokine-treated samples (Table S2A). Similarly, from a total of 41,201,153 and 48,903,611 reads in samples obtained from EVs from control and cytokine-treated islets, respectively, ∼58% and 53% of the total reads aligned to the reference genome. Approximately 28% of the aligned reads mapped to mature miRNAs in both experimental groups (6,588,292 reads in controls and 8,416,500 reads in cases, Table S2B).

Because samples were sequenced in two batches, principal component analysis (PCA) was performed to detect the presence of batch effects. PCA clustering using variance stabilized counts separated the samples into two batches, indicating the presence of batch effects in the dataset (Fig. S1A and S1C). A PCA plot clustering incorporating batch effect corrected variance stabilized normalized counts confirmed the removal of batch effects from the data of both islets (Fig. S1B) and EVs (Fig. S1D), as these samples clustered together without any separation based on batches.

Our methodology was robust as 1,110 and 890 miRNAs were retained in the datasets for control and cytokine-treated islets, respectively, after filtering for read counts. Yet, the evaluation was highly selective as a small, discrete subset of miRNAs were differentially expressed (DE) upon cytokine exposure. The volcano plot in Fig. 3A represents islet miRNAs that were significantly downregulated (blue dots) or upregulated (red dots) in cytokine-treated islets compared to control islets. Twenty DE miRNAs were identified in islets, of which 15 were up-regulated and 5 were down-regulated by cytokine treatment (Fig. 3B, Table S3-left panel). Similarly, the volcano plot in Fig. 4A depicts DE miRNAs identified from islet-derived EVs. Notably, we detected 14 DE miRNAs from EVs, 11 of which were up-regulated and 3 of which were down-regulated by cytokine treatment (Fig. 4B, Table S2-right panel).

**Figure 3.**
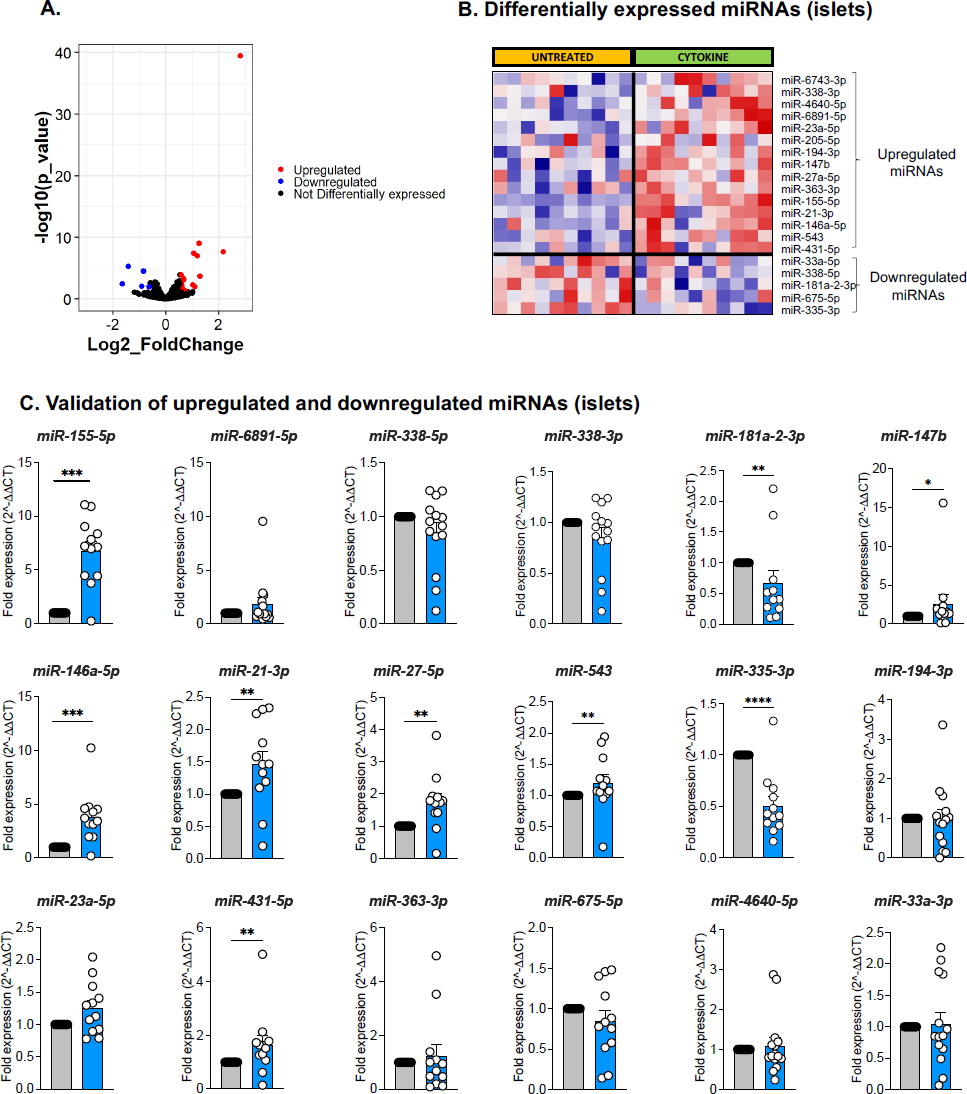
Identification and validation of differentially expressed islet-derived miRNAs. Human islets were treated with or without 50 U/mL IL-1β and 1000 U/mL IFN-γ for 24 hours. (A) Volcano plot showing upregulated and downregulated miRNAs from human islets treated with or without cytokines. (B) Heatmap illustrates the expression of the 20 differentially expressed miRNAs identified from small RNA sequencing analysis of control and cytokine-treated human islets (n=10; 5M/5F). (C) qRT-PCR validation of upregulated and downregulated miRNAs identified from sequencing analysis of human islets. Grey filled bars represent control and blue filled bars represent cytokine treatment. n=12; 7M/5F composition for C; *p <0.05, **p<0.01, ***p<0.001.

**Figure 4.**
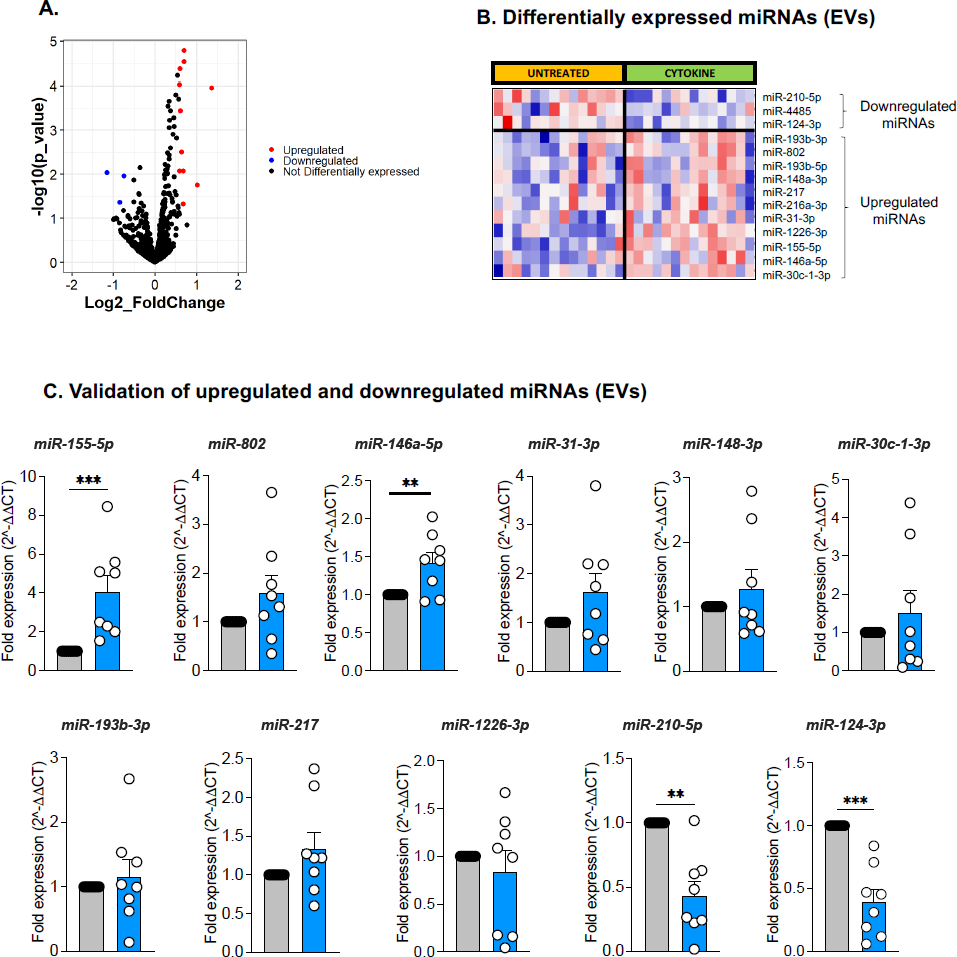
Identification and validation of differentially expressed islet-derived EV miRNAs. Human islets were treated with or without 50 U/mL IL-1β and 1000 U/mL IFN-γ for 24 hours and EVs were isolated from the culture supernatant. (A) Volcano plot showing upregulated and downregulated EV miRNAs isolated from human islets treated with or without cytokines. (B) Heatmap illustrates the expression pattern of 14 differentially expressed EV-miRNAs identified from small RNA sequencing analysis of isolated EVs from control and cytokine-treated islet culture supernatants. (N=14; 7M/7F). (C) qRT-PCR validation of upregulated and downregulated EV-miRNAs identified from small RNA sequencing analysis. Grey filled bars represent control and blue filled bars represent cytokine treatment EV data. n=8; 4M/4F for C; *p <0.05, **p<0.01, ***p<0.001.

### Validation of DE miRNAs from human islets and islet-derived EVs

Next, we validated the differential expression of miRNAs identified in our sequencing data using human islets from a separate set of cadaveric donors treated with or without IL-1β + IFN-γ. For this analysis, RNA was isolated and miRNA expression levels were measured by qRT-PCR. Of the 20 DE miRNAs in islets (Fig. 3B) and the 14 DE miRNAs in EVs (Fig. 4B), targeted analysis validated changes in 18 islet miRNAs (Fig. 3C, 13 up-regulated and 5 down-regulated) and 11 EV miRNAs (Fig. 4C, 9 up-regulated and 2 down-regulated). Therefore, ∼80% of the miRNAs identified in our small RNA sequencing showed similar expression patterns in qRT-PCR performed in a separate cohort of samples. However, only the changes in a subset of 7 up-regulated and 2 down-regulated islet miRNAs reached statistical significance (Fig. 3C). Analysis of EV miRNA expression patterns by qRT-PCR showed that 2 up-regulated and 2 down-regulated miRNAs were significantly changed (Fig. 4C). Collectively, the results from small RNA sequencing and qRT-PCR indicated that only two miRNAs (miR-155-5p and miR-146a-5p) were commonly upregulated in both islets and EVs upon cytokine treatment.

### Ranking of significant EV-derived miRNAs and pathway enrichment analysis

One goal of our analysis was to define a signature of EV miRNAs with potential to act as T1D biomarkers. To rank the discriminatory ability of DE EV-derived miRNAs to distinguish between cytokine and control conditions, we applied multiple machine learning algorithms. This analysis identified 10 miRNAs with the highest ranking based on coefficient values; a lower coefficient value indicated better performance across all algorithms (Fig. 5A). Among these 10 miRNAs, five miRNAs (miR-124-3p, miR-155-5p, miR-802, miR-30c-1-3p, and miR-146a-5p) were advanced for analysis in a clinical cohort. The criteria to advance these miRNAs was based on a combination of previous literature linking them with β cell dysfunction or autoimmunity, their rank in the integration of the machine learning analysis, and results from the qRT-PCR validation (Figs. 4C). The functional significance of these five miRNAs was identified using DIANA miRPath. Valid functional terms with an acceptable false discovery rate (FDR) of <0.05 are represented in Fig. 5B. The identified miRNAs were enriched for numerous terms associated with T1D, including “immune response”, “TLR signaling”, “inflammatory signaling”, “cell death”, and “apoptosis.”

**Figure 5.**
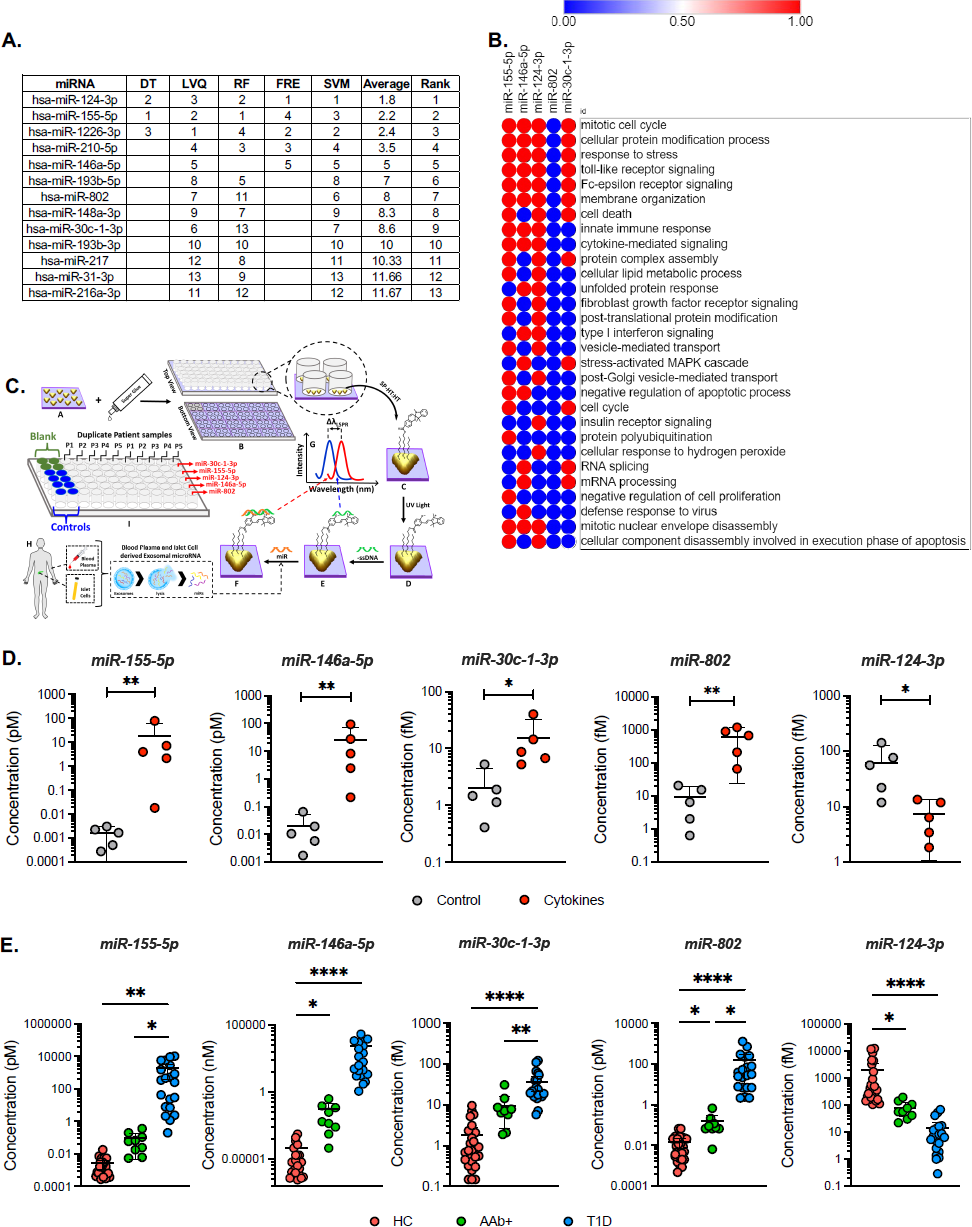
Differentially expressed miRNA rankings, miRNA regulated biological functions, and evaluation of miRNA expression in both human islet and plasma-derived EVs. (A) Table of the top differentially expressed miRNAs from islet-derived EV studies evaluated using machine learning tools to generate a coefficient value from the data that is used to rank the miRNA according to its signal performance (a lower value indicates better performance). (B) Predicted impact of miRNAs 155-5p, 146a-5p, 30c-1-3p, 802, and 124-3p upon biological processes. (C) Fabrication and development of custom, label-free LSPR-based biosensors and (D) LSPR-based biosensor determination of miRNA expression levels in EVs isolated from untreated and cytokine-treated human islet culture supernatant. (n=5; *p<0.05, **P<0.005). (E) Circulating plasma EVs were isolated from individuals with AAb+ but normoglycemic, new-onset T1D, and healthy controls. LSPR-based biosensor multi-plex miRNA assay showing the expression of the five miRNAs 155-5p, 146a-5p, 30c-1-3p, 802, and 124-3p from human plasma-derived EVs. (Healthy controls, n=26 (17M/9F); AAb+, n=8 (5M/3F); New-onset T1D, n=20 (11M/9F)). *p<0.05, **P<0.005, **P<0.0001.

### Validation of miRNA signatures in plasma samples of T1D clinical cohorts

To test the utility of this signature of miRNAs to act as biomarkers capable of identifying T1D risk, we isolated EVs from fasting plasma samples collected from individuals with AAb+ and recent onset Stage 3 T1D and age-matched non-diabetic controls. The clinical characteristics of these individuals are shown in Table 1. We designed custom, label-free localized surface plasmon resonance (LSPR)-based biosensors (Fig. 5C) to measure individual miRNA species. In brief, this method utilizes gold triangular nanoprisms that are functionalized with 75% spiropyran hexanethiol: 25% hexanthiol spacer (SP-HT:HT) to construct a biosensor that allows the measurement of a wide range of concentrations (e.g., nanomole to attomole) with high sensitivity and specificity using a simple UV-visible spectrophotometer in the absorbance mode (Fig. S2A-E). First, we validated our approach by using the LSPR-based biosensors to measure expression levels of miR-155-5p, miR-146a-5p, miR-30c-1-3p, miR-802 and miR-124-3p in EVs isolated from control and cytokine-treated human islets (Fig. 5D). The results of this analysis were concordant with results from qRT-PCR and RNA sequencing.

**Table 1:** Clinical and anthropometry characteristics of human plasma donors.

| **Disease Status** | **Total Subjects**<br>**(M/F)** | **Age**<br>**(Mean±SD)** | **BMI**<br>**(Mean±SD)** |
| --- | --- | --- | --- |
| Healthy controls | 26(17M/9F) | 11.38± 4.02 | 19.46±5.1 |
| AAb+ | 8(5M/3F) | 13.75±1.98 | 25.04±4.65 |
| New-onset T1D | 20(11M/9F) | 12.8±3.38 | 19.13±4.3 |

Next, we applied the biosensors in plasma-derived EVs from the clinical cohort. We observed a significant up-regulation of miR-155-5p, miR-146a-5p, miR-30c-1-3p, and miR-802, and a decreased abundance of miR-124-3p in plasma EVs from individuals with new-onset T1D compared to non-diabetic controls (Fig. 5E). In addition, we observed upregulation of miR-146a-5p and miR-802 and down-regulation of miR-124-3p in individuals with AAb+ (Fig. 5E) compared to non-diabetic controls.

In parallel, we assessed the expression of these five miRNAs in the clinical cohort using droplet digital (dd)PCR. Results from this analysis showed that both miR-155-5p and miR-146a-5p had significantly higher copy numbers in plasma EVs isolated from individuals with recent onset T1D compared to healthy controls (Fig. S3). Similar to data obtained from the LSPR-based biosensors, we observed a significant increase in the copy numbers of miR-146a-5p, miR-802 and miR-124-3p in AAb+ individuals compared to controls.

### Determination of localization and spatial distribution of miR-155-5p in human pancreatic tissues sections using *in situ* hybridization

While the content of miRNAs in EVs can inform biomarker strategies, intracellular miRNAs are known to regulate a variety of molecular pathways, including those associated with T1D pathophysiology. Additionally, it is now recognized that subcellular localization of miRNAs is associated with novel functions [23]. Here, we focused on miR-155-5p for downstream validation as this was the most highly up-regulated miRNA in our islet sequencing and qRT-PCR datasets (Fig. 3A-B). We determined both subcellular localization and expression levels of pre-miR-155-5p in intact pancreatic tissue sections from organ donors with AAb+, established T1D, and nondiabetic controls. smFISH imaging was performed on pancreatic tissue sections from human donors by co-labeling with DAPI for the nucleus and anti-insulin antibody to identify the β cells (Fig. 6). We developed an imaging-based bioinformatics approach to quantify the spatial distribution of pre-miR-155-5p specifically in single β cells.

**Figure 6.**
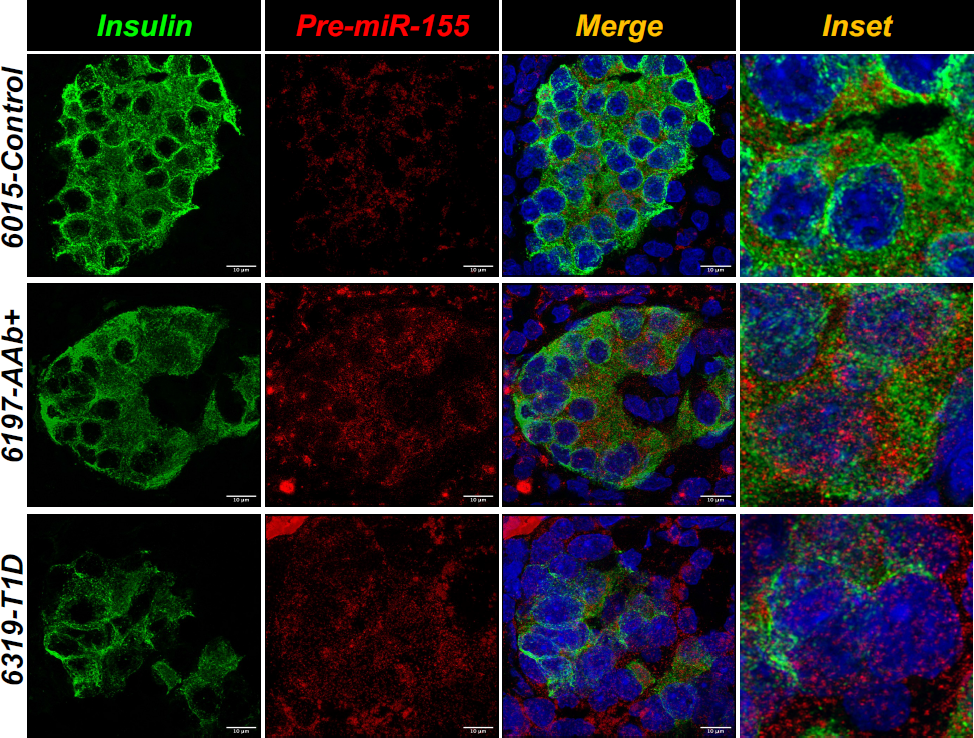
**In situ hybridization of pre-miRNA-155 expression in β cells of human pancreas tissue sections**. Representative images of single molecule in situ hybridization (smFISH) showing the expression of pre-miRNA-155 (red) in human pancreatic tissue sections obtained from non-diabetic control, AAb+, and T1D donors and counterstained with insulin (green) and nucleus (DAPI).

Next, we applied a supervised machine learning algorithm to our *in-situ* hybridization analyses. The spatial distribution of miRNA molecules in a single β cell were mathematically quantified by a series of features, which included the expression level of the pre-miRNA in the nucleus or cytoplasm (group 1), the location of single miRNAs in the β cell (group 2), the clustering of pre-miRNA-155-5p to itself (group 3), and the aggregation of pre-miRNA-155-5p at the nuclear/cellular boundary (group 4) (Fig. 7A-B). The group 1 and 2 features of smFISH images have been successfully leveraged in other studies to classify different cell types [24]. However, these two feature groups were not adequate to differentiate non-diabetic samples from the T1D samples with high accuracy. To this end, we integrated the mathematical model of clusters, i.e. Ripley’s K function and its derivatives [25,26], to characterize the degree of clustering for the miRNA. In addition, we also quantified the reciprocal clustering of pre-miRNA-155 as the group 4 feature.

**Figure 7.**
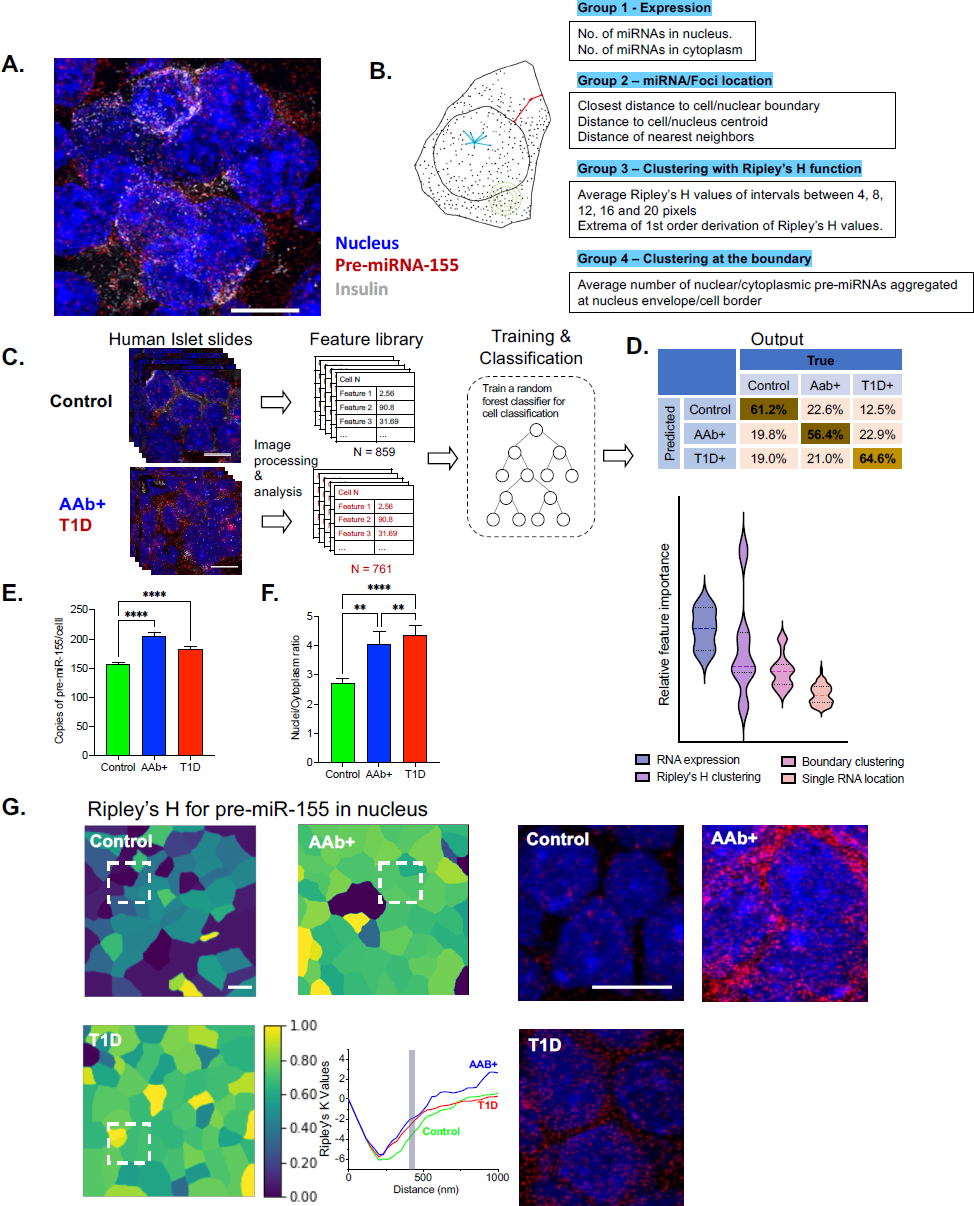
Machine learning-based spatial transcriptomics analysis of the pre-miRNA-155-expression in β cells. (A) Expression of the pre-miRNA-155 (red) was screened by smFISH imaging, with nucleus stained with DAPI (blue) and co-staining with insulin (gray). (B) Quantitative features that describe the location of the pre-miRNA molecule in a single β cell. Four groups of features are extracted for a single cell; this includes expression features (group 1), miRNA location features (group 2), clustering features (group 3), and boundary clustering features (group 4). (C) Supervised machine learning was applied to three categories of smFISH image data (control, T1D, and AAb+) for training and classification. (D) Single β cells belonging to different categories (control, T1D, and AAb+) were classified, the confusion matrix was produced, and distinctive features were ranked to show the expression and clustering of the pre-miRNAs that dominate the classification. (E) Spatial expression of total pre-miRNA-155 in each β cell was compared for all three categories (control, T1D, and AAb+). (F) The comparison of the nucleus/cytoplasm ratio of pre-miRNA-155 in β cells of control, T1D, and AAb+ samples. (G) Ripley’s H value at the distance of 457 nm (inset) for nuclear pre-miRNA-155 in a population of cells for all three categories (control, T1D, and AAb+), and the right panel indicates the real smFISH images of pre-miRNA-155 in the boxed area identified in the left. **p < 0.01, ****p < 0.0001.

The supervised machine learning algorithm was applied to 80% of total cells (N = 859 for Control, 307 for AAb+, and 454 for T1D) to establish the training network (Fig. 7C-D). Applying the algorithm to the remaining 20% of samples resulted in the classification of 61.2% of non-diabetic samples and 64.6% of T1D samples, and lower accuracy (56.4%) was observed with AAb+ sample prediction. Most importantly, this machine learning-based classification approach revealed a group of distinctive features of pre-miRNA-155 distribution between control, AAb+, and T1D samples. Interestingly, the spatial expression of the pre-miRNA and the local clustering effects dominated the classification process (Fig. 7C-D). In addition, spatial analysis of pre-miR-155 expression from insulin positive β cells showed an increase in pre-miR-155 copy numbers from individuals with AAb+ and T1D versus those of the control group (Fig. 7E). Moreover, sub-cellular localization of pre-miR-155 showed a spatial disparity in the pre-miR-155 expression, where we observed an increase in the nuclear to cytoplasmic ratio in the AAb+ and T1D samples compared to control donors (Fig. 7F). In line with this data, the color-coded cell images based on the Ripley’s K function at 457 nm suggest a different degree of pre-miRNA-155 clustering in tissue sections from control, AAb+, and T1D individuals, and this is clearly observed in the raw smFISH images (Fig. 7G). Our findings are the first to indicate that the physical distribution of pre-miRNA may be directly associated with β cell stress and dysfunction.

## Discussion

In this study, we aimed to identify islet miRNAs that may play a role in T1D pathogenesis and explore miRNA signatures of islet-derived EVs that could be leveraged as biomarkers to identify T1D risk. To this end, we utilized small RNA sequencing to identify miRNAs that were differentially expressed in both human islets and islet-derived EVs using an *in vitro* and organ-based model of T1D. We used a well-validated model of cytokine stress (50 U/mL of IL-1β and 1000 U/mL of IFN-γ) that has been shown to result in transcriptional changes that are correlated with those found in islets from persons with diabetes [27].

Several reports have shown that components of EV cargo such as nucleic acids (DNA and RNAs), lipids, and proteins are dynamically modulated in response to extrinsic and intrinsic cues and may reflect the cellular physiological or pathological state of a given cell type [28,29]. Moreover, EVs are protected from RNase-mediated degradation [30]. Hence, EV-derived miRNA signatures may serve as reliable markers to identify the health status of a cell. Methods to identify indicators of β cell stress can help dissect heterogeneity amongst at-risk individuals and could inform both the timing and selection of disease-modifying therapies. Current methods to do this in T1D are lacking.

Recently, we used total RNA sequencing to show that several types of RNA species, such as miRNAs, piRNAs, snoRNA, tRNAs, and poly(A) RNAs, are packaged and secreted through human islet-derived EVs [31]. However, the number of miRNAs identified by this approach was low. To address this lack of enrichment, here we utilized a robust small RNA sequencing approach from human islets and islet-derived EVs. This technique was robust, as we confidently identified 1110 miRNAs in islets and 890 miRNAs in EVs, which approached detection of ∼50% of the 2000 known human miRNAs. Interestingly, a much smaller subset of these miRNAs was differentially expressed between control and cytokine treatment (20 in islets and 15 in EVs), suggesting that miRNA biogenesis under stress conditions is engaged with a high degree of selectivity. Recent studies have shed light on the regulation of tissue retention and sorting of miRNAs into EVs. It has become clear that this process is highly regulated and depends on the specific sequence motifs carried by the miRNAs [32]. For instance, Liu et al. demonstrated that the sorting of miRNAs into EVs may occur through a process of liquid-liquid phase separation involving an RNA binding protein called YBX1 [33]. These findings have important implications for the use of EV-associated miRNAs as biomarkers, as the sorting process may affect the composition and abundance of miRNAs within EVs. In addition, understanding the mechanisms of miRNA sorting may lead to the development of new strategies for selectively loading miRNAs into EVs for therapeutic purposes. Further research is needed to fully elucidate the regulation of miRNA sorting into EVs and its implications for EV-based therapies and biomarker development.

Next, we applied a machine learning approach and multiple algorithms to prioritize a signature of miRNAs for clinical testing. Based on results from the machine learning analysis and validation of differentially expressed miRNAs using targeted qRT-PCR, we advanced miR-155-5p, miR-146a-5p, miR-30c-1-3p, miR-802, and miR-124-3p for analysis in plasma EVs isolated from children with AAb+, recent onset T1D, and healthy controls. This signature was informed also by published literature linking these miRNAs with inflammation and autoimmunity (miR-155-5p, miR-146a-5p, and miR-124-3p) [34,35]; acinar-ductal trans-differentiation in the pancreas (miR-802) [36]; glucose stimulated insulin secretion (miR124-3p) [37,38]; and fibrosis of the pancreas (miR-30c-1-3p) [39]. We used two methods to analyze this miRNA signature in the clinical samples. First, we employed a custom developed LSPR-based biosensing approach. Secondly, we used ddPCR, which has the ability to measure absolute copy numbers of any given miRNA. Overall, results from the LSPR assay and ddPCR analysis showed similar trends. However, the LSPR-based biosensor approach provided a better dynamic range and improved discriminatory ability across the groups. This approach is also quite robust as it does not require cDNA synthesis or PCR amplification steps. In addition, this biosensor works similarly to a microarray, which is more amenable to high-throughput analysis that might be required in a clinical study. Using this platform, we observed a significant upregulation of miR-155-5p, miR-146a-5p, miR-30c-1-3p, and miR-802-3p and a decreased abundance of miR-124-3p in plasma EVs from individuals with new-onset T1D compared to non-diabetic controls. In addition, miR-146a-5p and miR-802 were increased and miR-124-3p was decreased in plasma EVs from individuals with AAb+ compared to controls, suggesting that these miRNA signatures could reflect a phenomenon of both early and late β cell stress during T1D development.

In line with our data, a recent study by Januszewski et al. reported the association of 50 circulating plasma miRNAs with C-peptide levels in a cohort of individuals with and without T1D. They identified 16 miRNAs, including miR-146a and miR-155, that showed a strong association with plasma C-peptide levels [40]. Furthermore, this group found that 13 miRNAs, including miR-155 and miR-210, were strongly associated with detectable C-peptide in individuals with T1D [40]. Interestingly, Margaritis et al. performed a meta-analysis using published studies on T1D and identified seven miRNAs, including miR-181 and miR-210, that were significantly upregulated in individuals with T1D [41]. Consistent with these studies, our data identified upregulation of miR-146a and miR-155 in islets and islet-derived EVs, and we found increased levels of these miRNAs in our clinical cohort. In contrast, we observed a downregulation of miR-210 in islet-derived EVs (Figure 4), suggesting that source of miR-210 observed in other studies may emanate from another tissue type.

The final goal of our study was to identify ϕ3 cell miRNAs that play a role in T1D pathogenesis. Here, we aimed to test whether miR-155, the most highly upregulated miRNA across multiple of our datasets, was modulated similarly in human pancreases collected from individuals with AAb+ and T1D. miR-155 was a compelling target for translation into these tissues based on an abundance of previous literature linking it with T1D pathogenesis. Notably, miR-155 promotes the secretion of type I interferon by dendritic cells [42], stimulates an M1 phenotype in macrophages within the islet milieu [43,44], and plays a role in T-cell activation [45]. Previous studies in the NOD mouse model of T1D suggest that miR-155-5p can be transferred from β cells that undergo inflammatory insults to healthy β cells to amplify apoptosis [46] and from lymphocyte-derived EVs to β cells to enable disease development [20].

We found that pre-miR-155-5p expression was increased in ϕ3 cells from donors with AAb+ and T1D compared to non-diabetic donors. In this study, we also aimed to investigate the spatial localization of miRNAs within β cells of at-risk (AAb+) and early-onset T1D individual. Our results revealed that pre-miR-155-5p exhibits distinctive features of disease-predictive localization, including increased nuclear localization and clustering of miRNA signals. We utilized the high sensitivity of smFISH methodology along with machine learning algorithms for data analysis. The increased nuclear ratio of pre-miR-155 could be attributed to several possible mechanisms, such as nuclear processing of pre-miRNAs or nuclear import of processed miRNA from the cytosol through importin or nuclear pore complexes. Moreover, nuclear localization of pre-miR-155 could signify an increase in noncanonical activities of the miRNA, such as regulation of transcription via target gene promoters and cis-regulatory elements (CREs), regulation of chromatin structure (looping), and other epigenetic changes such as methylation, compaction of chromatin, and genomic remodeling. The self-clustering of miR-155 is suggestive of association with membrane-bound organelles or the formation of a scaffolding complex and a condensate of RNA [23,47]. These findings suggest that targeting miR-155 could be a promising approach for the treatment of T1D, and future studies could explore the potential of miR-155 inhibition as a therapeutic intervention in humans.

There are limitations of our study that should be noted. First, the sample size for the machine learning analysis to generate the EV miRNA signature was small. Typically, these algorithms perform most robustly with larger sample sizes and independent cohorts for both testing and validation. Despite this limitation, the predicted miRNAs performed well in distinguishing T1D and AAb+ individuals from non-diabetic controls in clinical testing. It is important to note that plasma-derived EVs contain a heterogeneous mix of EVs from different organs. While we identified these miRNAs from an islet-based stress model, we cannot presume they provide information solely about the health status of the β cell. Thus, it is possible that plasma levels of these miRNAs could be modulated in other inflammatory and/or autoimmune conditions. Whether this would diminish their utility in identifying T1D risk will require further testing.

Notwithstanding these limitations, we identified a robust panel of miRNAs that were modulated in individuals with T1D or AAb+. Moreover, the expression patterns observed in our cross-sectional cohorts suggest that miRNA expression levels may be modulated progressively during the evolution of T1D. Additional experiments should be performed in AAb+ cohorts followed longitudinally before and after seroconversion and through T1D development to confirm this hypothesis. Taken together, our results show that miRNA expression patterns in human pancreatic islets and islet-derived EVs change dynamically under inflammatory conditions and suggest that organ-based disease models can be leveraged to inform biomarker and therapeutic strategies for T1D.

## MATERIALS AND METHODS

### Experimental Design

The objective of this study was to investigate the expression of miRNAs in human islets and islet-derived EVs in response to pro-inflammatory cytokine treatment, and to determine the potential of circulating islet-derived EV miRNAs as a predictive diagnostic marker for early β cell stress or death. Islets from 10 cadaveric organ donors were treated with or without pro-inflammatory cytokines to model T1D in vitro. The differential expression of EV-derived miRNAs was analyzed using multiple machine learning classifications, and the top 10 miRNAs were ranked based on their predictive capability. We selected five miRNAs that have been shown to play a role in β cell health and function for further validation using a custom-developed label-free localized surface plasmon resonance (LSPR)-based biosensor approach. EV-miRNAs were isolated from islet-conditioned media and from plasma samples of individuals who are positive for islet autoantibody but normoglycemic, new-onset T1D, and healthy controls. Single-molecule in situ hybridization was used to determine the spatial expression of pre-miR-155 in pancreatic tissue sections. No power calculation was used for experiments involving human islets, and the sample number reflects the number of biological replicates used in each experiment. All samples for EV-miRNA measurement using LSPR were number-coded and blinded during processing and quantification.

### Culture of human islets

Human islets from 10 cadaveric donors were obtained from the Integrated Islet Distribution Program (IIDP). Islet donor and isolation characteristics, including donor age, sex, BMI, cause of death, measurements of islet purity and viability, ischemia duration, and culture time, are provided in Table S1 (17). Upon receipt, islets were placed in EV-depleted standard Prodo islet media (Prodo Labs, CA) and allowed to recover overnight. To investigate the effect of inflammation on islet miRNAs, islets were cultured with or without 50 U/mL IL-1β and 1000 U/mL IFN-γ for 24 hours. Untreated and cytokine-treated human islets and islet cell supernatants were harvested for downstream applications. For EV isolation, supernatants were collected and centrifuged for 30 minutes at 3000 x g to remove cellular debris, and EVs were then isolated using ExoQuick-TC (SBI Bioscience, USA), as per the manufacturer’s instructions.

### SEM and TEM imaging analysis of islets and EVs

Scanning electron microscopy (SEM) and transmission electron microscopy (TEM) imaging were performed at the Northwestern University Atomic and Nanoscale Characterization Experimental Center (NUANC) in Evanston, Illinois. Additional detail is provided in the Supplemental Methods.

### Nanoparticle tracking analysis

Nanoparticle tracking analysis (NTA; ParticleMetrix, Germany) was used to determine the size and concentration of EVs. Briefly, EVs isolated from human islet culture supernatants were diluted 1:3000 in ddH2O, and 1 mL of diluted sample was subjected to NTA. Results were analyzed using Zetaview Analysis software.

### Microarray chip-based tetraspanin staining

To determine the composition of EV cargo, we utilized ExoView microarray chips (Cat#EV-TETRA-C-T) printed with antibodies for the tetraspanin markers CD63, CD81, and CD9, and we performed the staining according to the directions provided by the manufacturer (ExoView, Cat# Cat#EV-TETRA-C-T). Additional detail is provided in the Supplemental Methods.

### Western blot analysis

EVs were washed twice with PBS and lysed with a buffer containing 50 mM Tris, 150 mM NaCl, 0.05% Deoxycholate, 0.1% IGEPAL, 0.1%SDS, 0.2% sarcosyl, 5% glycerol, 1 mM DTT, 1 mM EDTA, 10 mM NaF, and a cocktail of protease inhibitors (Roche) and phosphatase inhibitors (Roche). Protein concentrations were determined using the Lowry method (Bio-Rad), as per the manufacturer’s instructions. A total of 10 μL of protein was loaded onto 4-20% precast Tris-glycine gels (Bio-Rad), electroporated at 100 V, and protein transfer was performed using a PVDF membrane (Millipore). Blots were blocked with LI-COR PBS-blocking buffer and probed with the following primary antibodies at 4°C: anti-CD63 (1:1000 dilution; Cat#SC13118, Santa Cruz, mouse monoclonal, RRID: AB_627213) and anti-CD9 (1:1000 dilution; cat#SC5275, Santa Cruz, mouse monoclonal, RRID:AB_627877). Anti-rabbit or anti-mouse (1:10000) secondary antibodies were used for signal detection and band intensity was quantified using ImageStudio (LI-COR).

### RNA isolation from human islets and human islet-derived EVs

Total RNA was isolated from human islets using the miRNeasy Mini Kit (Qiagen, Germany), according to the manufacturer’s instructions. RNA quality and concentration were measured using 260/280 ratios (IMPLANT spectrophotometer, Germany), followed by assessment using the Agilent Bioanalyzer 2100 and small RNA assay chips (Agilent Technologies, USA). Exosomal RNA was isolated using the SeraMir RNA isolation kit (SBI Biosciences) per the manufacturer’s instructions. Briefly, isolated EVs were lysed with 350 µL SeraMir lysis buffer by vortexing for 15 seconds and allowing the lysate to stand at room temperature for 5 minutes. Next, 200 µL of 100% ethanol was added to the lysate and mixed by vortexing for 10 seconds. The lysate was transferred to a spin column and centrifuged at 13,000 rpm for 1 minute. The flow-through was discarded and the column was washed twice with 400 µL of wash buffer, followed by centrifugation at 13,000 rpm for 1 minute. Finally, the columns were dried by centrifugation at 13,000 rpm for 2 minutes, and RNA was eluted by adding 30 µL of elution buffer.

### Small RNA sequencing

Total RNA and miRNA were first evaluated using the Agilent Bioanalyzer for quantity, quality, and percent miRNA content in total RNA. The starting amount of RNA was 10-20 ng for each miRNA library. For miRNA library preparation, 0.5% of miRNA in total RNA was the cut-off to decide whether the sample should be enriched for miRNA. When needed, enrichment was performed following the small RNA library preparation procedure and the Ion Total RNA-Seq Kit v2 User Guide, Pub. No. 4476286 Rev. E (Life Technologies, USA). Each resulting barcoded library was quantified and assessed for quality using the Agilent Bioanalyzer. Multiple libraries were pooled in equal molarity. Eight µL of 100 pM pooled libraries were applied to the Ion Sphere Particles (ISP) template preparation and amplification was achieved using the Ion OneTouch 2, followed by ISP loading onto PI chips and sequencing using the Ion Proton semiconductor. Each PI chip is capable of generating approximately 50 million usable reads of 21-22 bp miRNA fragments. Sequence mapping was performed using Torrent Suite Software v4.6 (TSS 4.6). For details on miRNA sequence data analysis, see the Supplemental Methods.

### cDNA synthesis and quantitative RT-PCR for mRNA and miRNAs

To determine the expression of genes involved in EV biogenesis, 0.5-1 µg of total RNA was isolated from human islets and reverse transcribed using an M-MLV RT kit (Invitrogen, MA, USA). TaqMan primers (Applied Biosystems, CA, USA) were used for qRT-PCR. Β-actin was used as a normalizer to determine the expression of TP53, RAB27A, and RAB27B. The data were presented as fold expression using 2^-ΔΔ^Ct method compared to the untreated control.

For validation of DE miRNAs from human islets and islet-derived exosomal miRNAs, 1 µg of RNA isolated from human islets or 10 µL of exosomal RNA was reverse transcribed using the miRScript II kit (Qiagen, Germany), following manufacturer’s protocol. qRT-PCR was performed using miScript SYBR Green based PCR kit (Qiagen, Germany). miScript primer assays were used for the quantification of miRNA expression. All experiments were performed in duplicate. RNU6 and miR-484 served as internal reference genes for islet samples, and RNU6 alone served as the normalizer for EV samples (see Supplemental Methods for details). Relative fold expression against untreated control samples was calculated using the 2^-ΔΔ^Ct method [48].

### Feature selection and machine learning analysis to identify predictive miRNA signatures

We used the DE EV miRNAs from our small RNA sequencing to identify miRNA signatures with discriminatory ability, and we performed computational modeling using various machine learning algorithms. See Supplemental Methods for details.

### Preparation of human plasma-derived EVs and RNA isolation

Fasting plasma samples were collected from AAb+ children, pediatric subjects within 48 hours of diagnosis of T1D, and age-matched healthy control subjects [AAb+, n=8 (5M/3F); T1D, n=20 (11M/9F), and healthy controls, n=26 (17M/9F)]. The study was approved by the Indiana University School of Medicine Institutional Review Board. All study participants provided written informed consent or parental consent, and child assent was obtained in advance before any study participation, in accordance with the ethical guidance provided by Indiana University School of Medicine [49]. Citrated plasma samples were obtained by venipuncture or drawn from an existing intravenous line. Samples were processed and aliquoted within 1 hour of collection and then stored at −80°C until further use.

Plasma-derived EVs were isolated from plasma samples. Briefly, 500 µL of plasma sample was centrifuged at 1500 x g for 5 minutes to remove the residual cells and cellular debris. The supernatant was transferred to a fresh 1.5 mL tube, and 5 µL of de-fibrination reagent (Cat #TMEXO-1, System Biosciences, Palo Alto, CA) was added to the plasma to remove clotting factors prior to the isolation of EVs. The solution mixture was incubated at room temperature for 5 minutes with gentle mixing by flicking the tube. The sample was centrifuged at 10000 rpm for 5 minutes, and the supernatant was transferred to a clean tube. ExoQuick (Cat #EXOQ5A-1, System Biosciences, Palo Alto, CA) was added to the supernatant at a 1:4 ratio for 30 min at 4°C, followed by centrifugation at 1500 x g for 30 min. The supernatant was discarded without disturbing the pellet, and the isolated EVs were used for downstream applications.

### Validation of human plasma-derived EV miRNAs using ddPCR and LSPR-based biosensor

Isolated Evs were processed for total RNA isolation using the SeraMir Exosome RNA purification kit (Cat #RA808A-1, System Biosciences, Palo Alto, CA), according to the manufacturer’s instructions provided with the kit. RNA quality of each sample was measured using NanoDrop (Thermo Scientific, USA) and the Agilent Bioanalyzer Pico Assay using the Bioanalyzer 2100 Expert instrument (Agilent Technologies, Santa Clara, CA). cDNA was synthesized using the TaqMan MicroRNA Reverse Transcription kit (Cat# 4366596, Applied Biosystems, USA) and miRNA-specific RT primers. The expression of miRNAs was analyzed using ddPCR, as described previously [50,51], and a custom-developed LSPR-based biosensor (see Supplemental Methods). Synthetic mature miRNAs (Supplementary Table S4) were used as a positive control to gate the threshold, and the data were presented as copies/mL of plasma/serum EVs.

### BaseScope duplex detection assay

To define tissue expression patterns of pre-miR-155, single molecule fluorescent in-situ hybridization (smFISH) was performed using a BaseScope duplex detection assay, and machine-based learning was used to analyze images. See Supplemental Methods for details.

## QUANTIFICATION AND STATISTICAL ANALYSIS

To identify DE miRNAs, we performed paired sample analysis using the DESeq2 package in the R statistical program [52]. miRNAs with fold change (FC) ≥ 1.5 and p < 0.05 were considered as DE in cytokine-treated islets compared to untreated islets. For qRT-PCR analysis from islets and EVs we used Fishers exact-test to identify significant differences between cytokine-treated and untreated samples. Unpaired t-tests (Mann-Whitney test, non-parametric) were used for the analysis of biosensor data from islet-derived EV miRNAs with 95% confidence interval. A one-way ANOVA was used for the comparison of human plasma derived EV-miRNAs from healthy controls, AAb+, and new onset T1D. All the p values are represented as: * p ≤0.05, **p ≤0.001 (**), ***p ≤0.005 and ****p ≤0.0001.

## LIST OF SUPPLEMENTARY MATERIALS

### Supplemental Methods

Fig. S1: Principal component analysis on islets and EVs

Fig. S2: Calibration Curve generated using synthetic microRNAs

Fig. S3: Digital droplet PCR based determination of plasma derived EV-miRNAs

Fig. S4: UV-Visible absorption spectra and scanning electron microscope image of Au TNPs (cited in Supplemental Materials only)

Fig. S5: Representative UV-Vis extinction spectra showing Au TNP, SP-HT:HT, MC-HT:HT, - ssDNA-155-5p attachment and microRNA-155-5p hybridization (cited in Supplemental Materials only)

Table S1: Clinical and anthropometry characteristics

Table S2: Descriptive statistics of islets and EVs

Table S3: Differentially expressed miRNAs in islets and EVs

Table S4: ddPCR control gene synthesis

Table S5: Single stranded DNA (*-ssDNA)* oligomer sequences used in this study (cited in Supplemental Materials only)

Table S6: microRNA sequences that were used to generate calibration curves (cited in Supplemental Materials only)

Table S7: λ_LSPR_ responses of spiropyran based plasmonic nanosensors for microRNAs (cited in Supplemental Materials only)

Table S8: λ_LSPR_ responses of spiropyran based plasmonic nanosensors for nucleic acid ss-DNA (cited in Supplemental Materials only)

## Acknowledgments

This work was supported by NIH grants R01 DK093954 and DK127308 (to CEM), R01 DK060581 (to RGM) and U01DK127786 and UC4 DK 104166 (to RGM and CEM), VA Merit Award I01BX001733 (to CEM), 2-SRA-2019-834-S-B, JDRF 2-SRA-2018-493-A-B (to CEM and RGM), and gifts from the Sigma Beta Sorority, the Ball Brothers Foundation, and the George and Frances Ball Foundation (to CEM). F.S. was supported by JDRF postdoctoral fellowship (3-PDF-20016-199-A-N and JDRF 5-CDA-2022-1176-A-N). FS and JL were supported by NIH/NIDDK dkNET (U24DK097771). This research was performed with the support of the Network for Pancreatic Organ donors with Diabetes (nPOD; RRID:SCR_014641), a collaborative type 1 diabetes research project supported by JDRF (nPOD: 5-SRA-2018-557-Q-R), and The Leona M. & Harry B. Helmsley Charitable Trust (Grant#2018PG-T1D053, G-2108-04793). The content and views expressed are the responsibility of the authors and do not necessarily reflect the official view of nPOD. Organ Procurement Organizations (OPO) partnering with nPOD to provide research resources that are listed at http://www.jdrfnpod.org/for-partners/npod-partners. The authors acknowledge the support of the Islet and Physiology Core and the Translation Core of the Indiana Diabetes Research Center (P30DK097512). We thank Dr. Emily Anderson-Baucum for her assistance in editing the text of this manuscript.

## Author contributions

FS and CEM conceived and designed the study. FS, SRL, SH, RS, KY and AKL performed experiments and analysed the data. PK, JL, ST, XL, and GC performed computational analyses. FS and CEM interpreted the data and wrote the manuscript. RM provided materials and edited the manuscript. All authors provided critical revisions and edits to the manuscript. All authors read and approved of the final manuscript. CEM is the guarantor of this work, and both FS and CEM have verified the underlying data of this manuscript.

## Data and materials availability

The data from small RNA sequencing of human islets and islet-derived EVs were deposited in GEO database (accession number GSE160391). All the other data associated with this manuscript can be found in the manuscript or in the supplementary files.

## References and Notes

1. DiMeglio LA, Evans-Molina C, Oram RA. Type 1 diabetes. The Lancet. 2018 Jun 16;391(10138):2449–62.

2. Ziegler AG, Rewers M, Simell O, Simell T, Lempainen J, Steck A, Winkler C, Ilonen J, Veijola R, Knip M, Bonifacio E, Eisenbarth GS. Seroconversion to multiple islet autoantibodies and risk of progression to diabetes in children. JAMA. 2013 Jun 19;309(23):2473–9.

3. Insel RA, Dunne JL, Atkinson MA, Chiang JL, Dabelea D, Gottlieb PA, Greenbaum CJ, Herold KC, Krischer JP, Lernmark Å, Ratner RE, Rewers MJ, Schatz DA, Skyler JS, Sosenko JM, Ziegler AG. Staging presymptomatic type 1 diabetes: a scientific statement of JDRF, the Endocrine Society, and the American Diabetes Association. Diabetes Care. 2015 Oct;38(10):1964–74.

4. Herold KC, Gitelman SE, Ehlers MR, Gottlieb PA, Greenbaum CJ, Hagopian W, Boyle KD, Keyes-Elstein L, Aggarwal S, Phippard D, Sayre PH, McNamara J, Bluestone JA, AbATE Study Team. Teplizumab (anti-CD3 mAb) treatment preserves C-peptide responses in patients with new-onset type 1 diabetes in a randomized controlled trial: metabolic and immunologic features at baseline identify a subgroup of responders. Diabetes. 2013 Nov;62(11):3766–74.

5. Pescovitz MD, Greenbaum CJ, Krause-Steinrauf H, Becker DJ, Gitelman SE, Goland R, Gottlieb PA, Marks JB, McGee PF, Moran AM, Raskin P, Rodriguez H, Schatz DA, Wherrett D, Wilson DM, Lachin JM, Skyler JS, Type 1 Diabetes TrialNet Anti-CD20 Study Group. Rituximab, B-lymphocyte depletion, and preservation of beta-cell function. N Engl J Med. 2009 Nov 26;361(22):2143–52.

6. Orban T, Bundy B, Becker DJ, DiMeglio LA, Gitelman SE, Goland R, Gottlieb PA, Greenbaum CJ, Marks JB, Monzavi R, Moran A, Raskin P, Rodriguez H, Russell WE, Schatz D, Wherrett D, Wilson DM, Krischer JP, Skyler JS, Type 1 Diabetes TrialNet Abatacept Study Group. Co-stimulation modulation with abatacept in patients with recent-onset type 1 diabetes: a randomised, double-blind, placebo-controlled trial. Lancet. 2011 Jul 30;378(9789):412–9.

7. Rigby MR, DiMeglio LA, Rendell MS, Felner EI, Dostou JM, Gitelman SE, Patel CM, Griffin KJ, Tsalikian E, Gottlieb PA, Greenbaum CJ, Sherry NA, Moore WV, Monzavi R, Willi SM, Raskin P, Moran A, Russell WE, Pinckney A, Keyes-Elstein L, Howell M, Aggarwal S, Lim N, Phippard D, Nepom GT, McNamara J, Ehlers MR, T1DAL Study Team. Targeting of memory T cells with alefacept in new-onset type 1 diabetes (T1DAL study): 12 month results of a randomised, double-blind, placebo-controlled phase 2 trial. Lancet Diabetes Endocrinol. 2013 Dec;1(4):284–94.

8. Rigby MR, Harris KM, Pinckney A, DiMeglio LA, Rendell MS, Felner EI, Dostou JM, Gitelman SE, Griffin KJ, Tsalikian E, Gottlieb PA, Greenbaum CJ, Sherry NA, Moore WV, Monzavi R, Willi SM, Raskin P, Keyes-Elstein L, Long SA, Kanaparthi S, Lim N, Phippard D, Soppe CL, Fitzgibbon ML, McNamara J, Nepom GT, Ehlers MR. Alefacept provides sustained clinical and immunological effects in new-onset type 1 diabetes patients. J Clin Invest. 2015 Aug 3;125(8):3285–96.

9. Herold KC, Bundy BN, Long SA, Bluestone JA, DiMeglio LA, Dufort MJ, Gitelman SE, Gottlieb PA, Krischer JP, Linsley PS, Marks JB, Moore W, Moran A, Rodriguez H, Russell WE, Schatz D, Skyler JS, Tsalikian E, Wherrett DK, Ziegler AG, Greenbaum CJ, Type 1 Diabetes TrialNet Study Group. An Anti-CD3 Antibody, Teplizumab, in Relatives at Risk for Type 1 Diabetes. N Engl J Med. 2019 Aug 15;381(7):603–13.

10. Sims EK, Bundy BN, Stier K, Serti E, Lim N, Long SA, Geyer SM, Moran A, Greenbaum CJ, Evans-Molina C, Herold KC, Type 1 Diabetes TrialNet Study Group. Teplizumab improves and stabilizes beta cell function in antibody-positive high-risk individuals. Sci Transl Med. 2021 Mar 3;13(583):eabc8980.

11. Warshauer JT, Bluestone JA, Anderson MS. New Frontiers in the Treatment of Type 1 Diabetes. Cell Metabolism. 2020 Jan 7;31(1):46–61.

12. Bluestone JA, Buckner JH, Herold KC. Immunotherapy: Building a bridge to a cure for type 1 diabetes. Science. 2021 Jul 30;373(6554):510–6.

13. Grieco FA, Sebastiani G, Juan-Mateu J, Villate O, Marroqui L, Ladrière L, Tugay K, Regazzi R, Bugliani M, Marchetti P, Dotta F, Eizirik DL. MicroRNAs miR-23a-3p, miR-23b-3p, and miR-149-5p Regulate the Expression of Proapoptotic BH3-Only Proteins DP5 and PUMA in Human Pancreatic β-Cells. Diabetes. 2017 Jan;66(1):100–12.

14. Lakhter AJ, Pratt RE, Moore RE, Doucette KK, Maier BF, DiMeglio LA, Sims EK. Beta cell extracellular vesicle miR-21-5p cargo is increased in response to inflammatory cytokines and serves as a biomarker of type 1 diabetes. Diabetologia. 2018 May 1;61(5):1124–34.

15. Osmai M, Osmai Y, Bang-Berthelsen CH, Pallesen EMH, Vestergaard AL, Novotny GW, Pociot F, Mandrup-Poulsen T. MicroRNAs as regulators of beta-cell function and dysfunction. Diabetes/Metabolism Research and Reviews. 2016;32(4):334–49.

16. Sims EK, Lakhter AJ, Anderson-Baucum E, Kono T, Tong X, Evans-Molina C. MicroRNA 21 targets BCL2 mRNA to increase apoptosis in rat and human beta cells. Diabetologia. 2017 Jun;60(6):1057–65.

17. Carrillo-Lozano E, Sebastián-Valles F, Knott-Torcal C. Circulating microRNAs in Breast Milk and Their Potential Impact on the Infant. Nutrients. 2020 Oct;12(10):3066.

18. Alsaweed M, Lai CT, Hartmann PE, Geddes DT, Kakulas F. Human milk miRNAs primarily originate from the mammary gland resulting in unique miRNA profiles of fractionated milk. Sci Rep. 2016 Feb 8;6(1):20680.

19. Weber JA, Baxter DH, Zhang S, Huang DY, Huang KH, Lee MJ, Galas DJ, Wang K. The MicroRNA Spectrum in 12 Body Fluids. Clin Chem. 2010 Nov;56(11):1733–41.

20. Guay C, Kruit JK, Rome S, Menoud V, Mulder NL, Jurdzinski A, Mancarella F, Sebastiani G, Donda A, Gonzalez BJ, Jandus C, Bouzakri K, Pinget M, Boitard C, Romero P, Dotta F, Regazzi R. Lymphocyte-Derived Exosomal MicroRNAs Promote Pancreatic β Cell Death and May Contribute to Type 1 Diabetes Development. Cell Metabolism. 2019 Feb 5;29(2):348–361.e6.

21. Bashratyan R, Sheng H, Regn D, Rahman MJ, Dai YD. Insulinoma-released exosomes activate autoreactive marginal zone-like B cells that expand endogenously in prediabetic NOD mice. Eur J Immunol. 2013 Oct;43(10):2588–97.

22. Cianciaruso C, Phelps EA, Pasquier M, Hamelin R, Demurtas D, Ahmed MA, Piemonti L, Hirosue S, Swartz MA, Palma MD, Hubbell JA, Baekkeskov S. Primary Human and Rat β-Cells Release the Intracellular Autoantigens GAD65, IA-2, and Proinsulin in Exosomes Together With Cytokine-Induced Enhancers of Immunity. Diabetes. 2017 Feb 1;66(2):460–73.

23. Jie M, Feng T, Huang W, Zhang M, Feng Y, Jiang H, Wen Z. Subcellular Localization of miRNAs and Implications in Cellular Homeostasis. Genes. 2021 Jun;12(6):856.

24. Batish N, Stoeger T, Pelkmans L. Image-based transcriptomics in thousands of single human cells at single-molecule resolution. Nat Methods. 2013 Nov;10(11):1127–33.

25. Ripley BD. Modelling Spatial Patterns. Journal of the Royal Statistical Society: Series B (Methodological). 1977;39(2):172–92.

26. Kiskowski MA, Hancock JF, Kenworthy AK. On the use of Ripley’s K-function and its derivatives to analyze domain size. Biophys J. 2009 Aug 19;97(4):1095–103.

27. Eizirik DL, Sammeth M, Bouckenooghe T, Bottu G, Sisino G, Igoillo-Esteve M, Ortis F, Santin I, Colli ML, Barthson J, Bouwens L, Hughes L, Gregory L, Lunter G, Marselli L, Marchetti P, McCarthy MI, Cnop M. The Human Pancreatic Islet Transcriptome: Expression of Candidate Genes for Type 1 Diabetes and the Impact of Pro-Inflammatory Cytokines. PLOS Genetics. 2012 Mar 8;8(3):e1002552.

28. Grenier-Pleau I, Abraham SA. Extracellular vesicles tell all: How vesicle-mediated cellular communication shapes hematopoietic stem cell biology with increasing age. Experimental Hematology. 2021 Sep 1;101–102:7–15.

29. Kalluri R, LeBleu VS. The biology, function, and biomedical applications of exosomes. Science. 2020 Feb 7;367(6478):eaau6977.

30. Valadi H, Ekström K, Bossios A, Sjöstrand M, Lee JJ, Lötvall JO. Exosome-mediated transfer of mRNAs and microRNAs is a novel mechanism of genetic exchange between cells. Nat Cell Biol. 2007 Jun;9(6):654–9.

31. Krishnan P, Syed F, Jiyun Kang N, G. Mirmira R, Evans-Molina C. Profiling of RNAs from Human Islet-Derived Exosomes in a Model of Type 1 Diabetes. Int J Mol Sci [Internet]. 2019 Nov 25 [cited 2020 Apr 20];20(23). Available from: https://www.ncbi.nlm.nih.gov/pmc/articles/PMC6928620/

32. Garcia-Martin R, Wang G, Brandão BB, Zanotto TM, Shah S, Kumar Patel S, Schilling B, Kahn CR. MicroRNA sequence codes for small extracellular vesicle release and cellular retention. Nature. 2021 Dec 22;1–6.

33. Liu XM, Ma L, Schekman R. Selective sorting of microRNAs into exosomes by phase-separated YBX1 condensates. Pfeffer SR, Zhao YG, editors. eLife. 2021 Nov 12;10:e71982.

34. Jiang Z, Yang M, Jin J, Song Z, Li C, Zhu Y, Tang Y, Ni C. miR-124-3p Down-Regulation Influences Pancreatic-β-Cell Function by Targeting Secreted Frizzled-Related Protein 5 (SFRP5) in Diabetes Mellitus. Journal of Biomaterials and Tissue Engineering. 2021 Apr 1;11(4):711–7.

35. Assmann TS, Duarte GCK, Brondani LA, de Freitas PHO, Martins ÉM, Canani LH, Crispim D. Polymorphisms in genes encoding miR-155 and miR-146a are associated with protection to type 1 diabetes mellitus. Acta Diabetol. 2017 May 1;54(5):433–41.

36. Ge W, Goga A, He Y, Silva PN, Hirt CK, Herrmanns K, Guccini I, Godbersen S, Schwank G, Stoffel M. miR-802 Suppresses Acinar-to-Ductal Reprogramming During Early Pancreatitis and Pancreatic Carcinogenesis. Gastroenterology. 2022 Jan 1;162(1):269–84.

37. Sebastiani G, Po A, Miele E, Ventriglia G, Ceccarelli E, Bugliani M, Marselli L, Marchetti P, Gulino A, Ferretti E, Dotta F. MicroRNA-124a is hyperexpressed in type 2 diabetic human pancreatic islets and negatively regulates insulin secretion. Acta Diabetol. 2015 Jun;52(3):523–30.

38. Plaisance V, Waeber G, Regazzi R, Abderrahmani A. Role of MicroRNAs in Islet Beta-Cell Compensation and Failure during Diabetes. J Diabetes Res. 2014;2014:618652.

39. Wu M, Liang G, Duan H, Yang X, Qin G, Sang N. Synergistic effects of sulfur dioxide and polycyclic aromatic hydrocarbons on pulmonary pro-fibrosis via mir-30c-1-3p/ transforming growth factor β type II receptor axis. Chemosphere. 2019 Mar;219:268–76.

40. Januszewski AS, Cho YH, Joglekar MV, Farr RJ, Scott ES, Wong WKM, Carroll LM, Loh YW, Benitez-Aguirre PZ, Keech AC, O’Neal DN, Craig ME, Hardikar AA, Donaghue KC, Jenkins AJ. Insulin micro-secretion in Type 1 diabetes and related microRNA profiles. Sci Rep. 2021 Jun 3;11(1):11727.

41. Margaritis K, Margioula-Siarkou G, Margioula-Siarkou C, Petousis S, Kotanidou EP, Christoforidis A, Pavlou E, Galli-Tsinopoulou A. Circulating serum and plasma levels of micro-RNA in type-1 diabetes in children and adolescents: A systematic review and meta-analysis. European Journal of Clinical Investigation. 2021;51(7):e13510.

42. Zhou H, Huang X, Cui H, Luo X, Tang Y, Chen S, Wu L, Shen N. miR-155 and its star-form partner miR-155* cooperatively regulate type I interferon production by human plasmacytoid dendritic cells. Blood. 2010 Dec 23;116(26):5885–94.

43. Vergadi E, Ieronymaki E, Lyroni K, Vaporidi K, Tsatsanis C. Akt Signaling Pathway in Macrophage Activation and M1/M2 Polarization. The Journal of Immunology. 2017 Feb 1;198(3):1006–14.

44. Arranz A, Doxaki C, Vergadi E, Martinez de la Torre Y, Vaporidi K, Lagoudaki ED, Ieronymaki E, Androulidaki A, Venihaki M, Margioris AN, Stathopoulos EN, Tsichlis PN, Tsatsanis C. Akt1 and Akt2 protein kinases differentially contribute to macrophage polarization. Proceedings of the National Academy of Sciences. 2012 Jun 12;109(24):9517–22.

45. Tanaka PP, Oliveira EH, Vieira-Machado MC, Duarte MJ, Assis AF, Bombonato-Prado KF, Passos GA. miR-155 exerts posttranscriptional control of autoimmune regulator (Aire) and tissue-restricted antigen genes in medullary thymic epithelial cells. BMC Genomics. 2022 May 28;23(1):404.

46. Roggli E, Britan A, Gattesco S, Lin-Marq N, Abderrahmani A, Meda P, Regazzi R. Involvement of microRNAs in the cytotoxic effects exerted by proinflammatory cytokines on pancreatic beta-cells. Diabetes. 2010 Apr;59(4):978–86.

47. Suzuki HI, Onimaru K. Biomolecular condensates in cancer biology. Cancer Sci. 2022 Feb;113(2):382–91.

48. Livak KJ, Schmittgen TD. Analysis of relative gene expression data using real-time quantitative PCR and the 2(-Delta Delta C(T)) Method. Methods. 2001 Dec;25(4):402–8.

49. Ocaña GJ, Sims EK, Watkins RA, Ragg S, Mather KJ, Oram RA, Mirmira RG, DiMeglio LA, Blum JS, Evans-Molina C. Analysis of serum Hsp90 as a potential biomarker of β cell autoimmunity in type 1 diabetes. Picard D, editor. PLoS ONE. 2019 Jan 10;14(1):e0208456.

50. Das Gupta S, Ndode-Ekane XE, Puhakka N, Pitkänen A. Droplet digital polymerase chain reaction-based quantification of circulating microRNAs using small RNA concentration normalization. Scientific Reports. 2020 Jun 2;10(1):9012.

51. Zhao G, Jiang T, Liu Y, Huai G, Lan C, Li G, Jia G, Wang K, Yang M. Droplet digital PCR-based circulating microRNA detection serve as a promising diagnostic method for gastric cancer. BMC Cancer [Internet]. 2018 Jun 22 [cited 2020 Sep 5];18. Available from: https://www.ncbi.nlm.nih.gov/pmc/articles/PMC6013872/

52. Love MI, Huber W, Anders S. Moderated estimation of fold change and dispersion for RNA-seq data with DESeq2. Genome Biology. 2014 Dec 5;15(12):550.

