## Supplemental Methods and Figures for "Title: β Cell microRNAs Function as Molecular Hubs of Type 1 Diabetes Pathogenesis and as Biomarkers of Diabetes Risk"

##### SEM and TEM imaging analysis of islets and EVs

Scanning electron microscopy (SEM) and transmission electron microscopy (TEM) imaging were performed at the Northwestern University Atomic and Nanoscale Characterization Experimental Center (NUANC) in Evanston, Illinois. After indicated treatments, islets were fixed in freshly prepared buffer containing 2.5% glutaraldehyde, 2% paraformaldehyde, and cacodylate buffered fixative and then stored at 4°C. On the following day, the samples were exchanged into fresh fixatives. The samples were microwave processed in a Pelco Biowave built-in with a cold spot and vacuum to enhance the sample processing. Samples were washed thrice with cacodylate buffer and incubated in a solution containing 1% OsO4 and 0.8% $KFeCN_6$. Subsequently, the samples were washed twice with cacodylate buffer and two washes with diH2O. The samples were then transferred to a solution containing 1% uranyl acetate (UA) in diH2O and washed thrice with diH2O. Samples were dehydrated twice with a series of different ethanol concentrations: 30%, 50%, 70%, 90%, and 100%. During the 100% ethanol step, the samples were divided and designated for SEM- or TEM-specific processing. For SEM imaging, the samples were dried using a Tousimis Samdri-795 Semi-Automated Critical Point Dryer. Dried samples were transferred into a SEM stub and coated with Pt/Pd. Imaging was performed on a Hitachi SU8030 SEM using an accelerating 2 or 5 kV voltage. Samples for TEM imaging were processed in a transitional fluid of acetone infiltrated with EMBed812 resin and polymerized at 60°C for 24 hours. Ultrathin sections were cut using a Diatome diamond knife using a Leica Ultracuts ultramicrotome. Cross-sections with a thickness of 90 nm were made and collected onto copper grids and imaged in a JEOL 1230 TEM with an 80kV accelerating voltage using a Gatan Orius camera.

**Microarray Chip-based tetraspanin staining**

The antibody coated microarray chips were incubated at room temperature for 15 minutes and pre-scanned using the ExoView R200 automated imaging system (NanoView Bioscience, USA). Chips were transferred to 24 well plates using tweezers, and EVs were isolated from control and cytokine-treated human islet culture supernatant using in-house developed dual chromatography-based approach. Isolated EVs were diluted (1:10) using 1X sample incubation buffer, and 50 µL of the diluted sample was added to the chip and incubated for 16-24 hours at room temperature. After the incubation period, the chips in 24 well plates were transferred to an automated plate washer and washed thrice with 1000 µL of solution A, then 750 µL of buffer was removed and incubated with 250 µL fluorescently labeled capture antibodies: anti-CD9 (CF®488A), anti-CD81 (CF®555), and anti-CD63 (CF®647) at 1:500 dilution in blocking solution II and incubated for 1 hr in an orbital shaker at 500 rpm. Next, the chips were washed thrice with 1X solution A, thrice with 1X solution B, and then thrice with DI water. The chips were placed at 45° angle, slowly pulling the chip out of the water, and placed on an absorbent paper and protected from light. The chips were imaged using an ExoView R200 automated imager, and the data were analyzed using ExoView data analysis software.

**miRNA sequence data analysis**

The raw sequencing files were aligned to the human genome (hg19) using STAR aligner v2.5.3a. Mature miRNAs were annotated using miRBase v20. miRNAs that had a total of less than 10 read counts were removed from further analysis. Read alignment, annotation, and generation of raw counts were completed using Partek Flow v6.0.18.0110 (Copyright ©; 2018 Partek Inc., St. Louis, MO, USA). Paired sample analysis was used to identify differentially expressed (DE) miRNAs using the DESeq2 package in the R statistical program (1). miRNAs with fold change (FC) $\geq 1.5$ and $p < 0.05$ were considered as DE in cytokine-treated islets (cases) compared to untreated islets (controls).

##### Selection of internal reference genes for quantitative RT-PCR for miRNAs

Potential reference miRNAs capable of serving as normalization strategies for qRT-PCR were identified using sequencing data from human islets and islet-derived EVs. A molecule was considered a potential reference gene if expressed in high amounts in both cases and controls; therefore, only miRNAs with at least 10 read counts in all the samples were considered. Further, any miRNA which showed a statistical significance of $p < 0.1$ between cases and controls was removed. To assess the relative stability of miRNAs, we calculated the co-efficient of variations (CVs), and miRNAs with a CV < 0.3 were retained. In addition, miRNAs falling within the 10th percentile and showing a normal distribution with normality p-value > 0.6 (Shapiro-Wilk test) were identified. As a final filtering criterion, miRNAs with a fold change < 1.05 were considered potential reference genes. miRNAs identified from islets and islet-derived EVs were tested using qRT-PCR. RefFinder (http://150.216.56.64/referencegene.php#) was used to determine the most stable miRNA as it compares the relative stability across different methods such as delta Ct, BestKeeper, NormFinder, and GeNorm and gives a comprehensive ranking of the input RNAs.

##### Feature selection and machine learning analysis to identify predictive miRNA signatures

For feature selection, we used Learning Vector Quantification (LVQ) implemented in the ‘caret’ R package to compute the importance values of features and any feature above a 0.7 importance value was considered significant and selected for machine learning analysis. Following variable selection, several supervised machine learning classifiers, including Decision Tree, Random Forest, and Support Vector Machine (SVM), were used to predict cytokine-treated islet-derived EVs from untreated islet-derived EVs. The R packages ‘rpart’, ‘randomforest’, and ‘e1701’ were used for the decision tree, random forest, and SVM analysis, respectively. In the random forest analysis, the number of trees was set to 500.

**Functional prediction of validated miRNAs**

Functional enrichment analysis of the five selected miRNAs was carried out using DIANA miRPath v.3 (2). Specifically, TarBase was selected for target prediction, followed by gene ontology analysis. Biological process terms with FDR <0.05 were considered to be significant. Common and unique functional terms between all five miRNAs were identified.

**LSPR-based biosensor construction**

6-mercapto-1-hexanol was obtained from TCI Chemicals. RNase-free sterile water was purchased from Baxter Healthcare Corporation. RBS 35 Detergent, 18x18 mm glass coverslips, trifluoroacetic acid (TFA, 99%), and dichloromethane (DCM) were purchased from Fisher Scientific. Chloro(triethylphosphine) gold (I) ($Et_3PAuCl$, 97%) was purchased from Gelest Inc. Poly(methylhydrosiloxane) (PMHS, $M_n$ = 1700-3300), triethylamine (TEA, 98%), hexamethylene tetramine (HMTA, ≥99.0%), 1,3,3-trimethyl-2-methylene indoline (TMMI, 97%), dimethyl aminopyridine (DMAP, ≥98.0%), 1-hexanethiol (98%), and ACS grade acetonitrile ($CH_3CN$, 99.9%) were purchased from Sigma-Aldrich. 4-hydroxy benzoic acid (BA, 99%) and (N,N-dicyclohexylcarbdiimide (99%) were purchased from Acros Organics. 3-Mercaptopropyl-trimethoxysilane (MPTMS, 94%) was purchased from Alfa Aesar. Ethanol (200 proof) (EtOH) was purchased from Decon Labs. Ethyl acetate (EtOAc) and Methanol (MeOH) were purchased from Pharmco. No-bottom 96-multiwell plates were purchased from Greiner Bio-one. Krazy glue was obtained from Office Depot. All water was purified using a Thermo Scientific Barnstead Nanopure system. All single-stranded oligonucleotide (Table S5) for 155-5p (-ssDNA-155-5p), 802 (-ssDNA-802), 30c-1-3p (-ssDNA-30c-1-3p),124-3p (-ssDNA-124-3p), 146a-5p (- ssDNA-146a) and mature microRNAs (Table S6), microRNA-155-5p, microRNA-802, microRNA-124-3p, microRNA-30c-1-3p and microRNA-146a-5p, were synthesized by Integrated DNA Technologies (IDT). The single-stranded oligonucleotide, mature microRNAs, and patient samples were stored at -80° C. PBS buffer (pH = 7.2) was prepared using RNase-free sterile water. Ethanol and acetonitrile were purged with $N_2$ for 30 min prior to use.

<u>Silanization of Glass Coverslips</u>: Glass coverslips with 18x18 mm dimensions were functionalized with MPTMS according to our previously published procedure (3–8). Briefly, glass coverslips were incubated in 10% (v/v) aqueous RBS detergent solution at 90° C and were sonicated for 15 min. Coverslips were then rinsed several times with nanopure water and incubated in a 1:1 (v/v) hydrochloric acid:methanol solution for 30 min at room temperature. Coverslips were then thoroughly rinsed with nanopure water, followed by drying in a vacuum oven at 60° C for at least 12 hrs. Next, coverslips were brought to room temperature, followed by incubation in a 15% (v/v) solution of MPTMS in $N_2$ purged ethanol for 30 min. The coverslips were then sonicated at least three times in ethanol for 15 min each for washing. After completion of the ethanol washes, the coverslips are dried in a vacuum oven at 120° C for at least 3 hrs. Finally, the MPTMS-functionalized coverslips were stored at 4° C for up to one week.

Synthesis of Gold Triangular Nanoprisms (Au TNPs): Au TNPs were chemically synthesized according to our previously published procedure with minor modifications (4–9). Briefly, gold salt, $Et_3PAu(I)Cl$ (18.4 mg, 0.05 mmol), was dissolved in 40 mL $CH_3CN$ and stirred for 10 min at room temperature. Next, 38.0 μL (0.273 mmol) of TEA was added to the gold salt solution and heated to 38° C. At this point, 0.6 mL of PMHS was added, and the reaction was allowed to proceed with gentle stirring. During the reaction, the color of the solution changed from colorless to pink to light purple or blue. Once the solution displayed a stable LSPR dipole peak ($\lambda_{LSPR}$) position at 800 nm in $CH_3CN$, the reaction was stopped by removing the container from the hot plate. The solution was immediately centrifuged at 7500 rpm for 10 sec, and the dark purple color solution was transferred to previously prepared MPTMS-functionalized 18x18 coverslips and incubated for exactly 1 hr, followed by rinsed with a copious amount of $CH_3CN$, dried under $N_2$ flow, and then stored under $N_2$ at 4° C for further use. Au TNP-bound glass coverslips were glued to a no-bottom 96-multiwell plate. This was accomplished by applying a small amount of super glue to the edge of 4 wells on the bottom side of the 96 well plate before placing a coverslip over the wells and gently pressing for 10 sec. Plates were allowed to dry for at least 2 hr at room temperature prior to use.

Synthesis of Receptor Binding Motif, SP-HT: The synthesis of spiropyran hexanethiol (SP-HT) was conducted using our published procedure with modifications (10). The synthesis consisted of three major steps: (1) synthesis of 3-formyl-4-hydroxybenzoic acid (BA-COOH), (2) synthesis of spiropyran carboxylic acid (SP-COOH), and (3) synthesis of spiropyran hexanethiol (SP-HT). (1) Synthesis of BA-COOH: BA-COOH was synthesized utilizing a Duff reaction. 5.0 g (36.2 mmol) of 4-hydroxy-benzoic acid (BA) was added to a 100 mL 2-neck round bottom flask and mixed with 15 mL of trifluoroacetic acid (TFA) under nitrogen with stirring for 30 min at room temperature. Separately, 5 g (36.2 mmol) of hexamethylenetetramine (HMTA) was mixed with 15 mL of TFA. The mixture was then added dropwise to the round bottom flask containing the BA. After completion of the addition, the reaction vessel was transferred to a preheated 250$^{o}$ C oil bath and refluxed for 3 hr under $N_2$. ESI analysis was conducted to confirm BA was fully reacted by the disappearance of m/z 138. BA-COOH was then precipitated using 4 N hydrochloric acid (HCl) over 3 hr with vigorous stirring and recovered using vacuum filtration. The product was rinsed thoroughly with nanopure water and dried under vacuum overnight. ESI-MS analysis was used to confirm the presence of BA-COOH, MS (ESI): *m/z* = 166. <u>(2) Synthesis of SP-COOH:</u> 2.5 g (15.1 mmol) of synthesized BA-COOH was added to a 100 mL 2-neck round bottom flask under $N_2$. 40 mL of purged ethanol was added and 3.8 mL (21.8 mmol) of 1,3,3-trimethyl-2-methylene indoline (TMMI) was obtained and added under $N_2$ with stirring. The reaction mixture was then placed in an oil bath and brought to reflux at 175° C for 3 hr. An ESI-MS analysis was conducted to determine that BA-COOH was no longer present by the disappearance of m/z 166. SP-COOH was purified using a silica gel column with a solvent gradient of dichloromethane (DCM) and methanol (MeOH). The SP-COOH product was obtained at 5 % MeOH and the remainder at 10 % MeOH. The fractions were concentrated using a rotatory evaporator (rotovap) and dried under a vacuum overnight. An ESI-MS product analysis was conducted to confirm the presence of SP-COOH, MS (ESI): *m/z* = 320. <u>(3) Synthesis of spiropyran hexanethiol (SP-HT)</u>: This process utilized a DCC-NHS coupling of SP-COOH with 6-mercapto-1-hexanol to produce the final product, SP-HT. First, 5 g (15.7 mmol) of SP-COOH was added to 100 mL of purged dichloromethane (DCM) in a 250 mL 2-neck round bottom flask under nitrogen, and 2.5 g (18.8 mmol) of 6-mercapto-1-hexanol and 0.21 g (1.7 mmol) of dimethyl amino pyridine (DMAP) were both added directly to the reaction mixture. The reaction was stirred under $N_2$ in an ice bath until the internal temperature reached 0$^{o}$ C. Separately, 3.9 g (18.8 mmol) of N,N-dicyclohexylcarbdiimide (DCC) was dissolved in 100 mL of purged DCM. The DCC solution was added to the reaction flask dropwise over 40 min. The reaction then slowly reached room temperature with stirring and was allowed to react overnight under $N_2$. Before the reaction was stopped, an ESI analysis was conducted to determine that SP-COOH was no longer present by the disappearance of m/z 320. The reaction solution was then concentrated under rotovap and purified with a silica gel column. The column utilized a hexane (HEX) and ethyl acetate (EtOAc) gradient to obtain the desired SP-HT product. SP-HT was obtained at 40-50% EtOAc, and a 100% MeOH wash of the column was used to remove any remaining product. The fractions were concentrated using a rotovap and then dried under a vacuum overnight. An ESI-MS analysis of the product was conducted to confirm the presence and purity of SP-HT by the appearance of molecular weight *m/z* = 437, MS (ESI).

<u>Fabrication of LSPR-Based Biosensors:</u> Following SP-HT synthesis, a mixture of 75% SP-HT (0.025M) and 25% hexanethiol (HT) (0.025M) in acetonitrile (SP-HT:HT) was prepared. This solution was then poured into previously constructed Au TNP-containing 96 well plate where each well was filled with ~200 µL of SP-HT:HT solution and then incubated overnight. The wells were then rinsed with copious amounts of $CH_3CN$ and dried with $N_2$ flow. The formation of mixed self-assembled monolayers (SAMs) of SP-HT:HT was confirmed by measuring the $\lambda_{LSPR}$. Utilizing the photochromic nature of SP, the 96 well plates were then irradiated for ~5 min with a Kessil photoreaction light (KSPR160L-370) at 50% power. This process allowed the closed-ring spiropyran (SP) structure to be converted into the open-ring, zwitterionic merocyanine (MC) form. $\lambda_{LSPR}$ was used to confirm this photoisomerization. Once in the activated MC form, the now fully

functional zwitterionic state could be utilized to bind the target receptor of choice, 300 μL of 10 μM -ssDNA-X (where X = -155-5p, -146a-5p, 802, -124-3p or -30c-1-3p) via a charge-charge interaction, as described below. The -ssDNA-X bound MC is considered as LSPR-based biosensor.

<u>Spectroscopy and Microscopy Characterizations</u>: Absorption and extinction spectra in the range of 700-1000 nm were collected with a SpectraMax M5 microplate reader from Molecular Devices, LLC. All absorption spectra were collected in PBS buffer (pH 7.2) to keep the bulk refractive index constant. A blank glass coverslip immersed in PBS buffer was used as a background, and an LSPR-based biosensor incubated in PBS buffer only was considered the blank reference. The chemically synthesized Au TNPs attached onto the MPTES-functionalized glass coverslips inside the 96 multiwell plate wells were characterized using a JEOL 7800F scanning electron microscopy (SEM) (Fig. S4).

<u>Development of Biomarker Calibration Plots</u>: The LSPR extinction spectra of the LSPR-based biosensors were collected in PBS buffer to determine their dipole peak position ($\lambda_{LSPR}$) before and after analyte adsorption. Specifically constructed biosensors were incubated overnight in 300 μL of microRNA-X, where X = -155-5p, -146a-5p, -802, -124-3p, or -30c-1-3p, solution of different concentrations (range of 100 nM to 100 aM) in 10% human plasma/PBS buffer solution (prepared via serial dilution) (Table S7). Each analyte-bound biosensor was washed with PBS buffer to remove any non-specifically adsorbed biomolecules, the LSPR extinction spectra were collected, and the $\lambda_{LSPR}$ was determined. The difference between the $\lambda_{LSPR}$ before and after analyte adsorption ($\Delta\lambda_{LSPR}$) was used to develop calibration plots. The false positive analysis was conducted by incubating the LSPR-based biosensor with receptor molecule functionalization in a PBS buffer

solution without any analytes present. False negative analysis was conducted by incubating the biosensors without any receptor molecules present in the 10.0 nM biomarker solution (Fig. S5).

Analyzing Clinical Samples: Approximately 10 μL of human plasma-derived EV miRNA samples were diluted in 3 mL total of PBS buffer (pH 7.2), and then 300 μL were transferred to individual wells acting as a LSPR-based biosensor. Each biomarker was assayed in duplicate measurements. For islet cell-derived exosomal microRNA samples, we followed the same procedure. LSPR-based biosensors were incubated into microRNA samples for overnight, followed by rinsing with excess nanopure water, and finally LSPR extinction spectra were collected. The $\lambda_{LSPR}$ was determined for each well, and the difference in the $\lambda_{LSPR}$ from before (meaning after receptor -ssDNA-X attachment) and after patient sample microRNA adsorption to LSPR-based biosensor was determined ($\Delta\lambda_{LSPR}$) (Table S8). The $\Delta\lambda_{LSPR}$ value was then plugged into the biomarker calibration plots to quantify the miRNA concentration for each patient sample as described below.

Data Processing and Statistical Analysis:

*Processing UV-Vis Extinction Spectra:* From the UV-Visible extinction spectra, $\lambda_{LSPR}$ was determined through curve fitting using Origin software, and $\Delta\lambda_{LSPR}$ was derived by taking the difference between the $\lambda_{LSPR}$ of the biosensors before and after attachment of the target miRNA analyte.

*Processing Calibration Curves, Limit of Detections, and Concentration of Target miRNAs in Patient Plasma and Islet Cell Derived EV lysate:* As shown in Fig S2, calibration curves were developed by plotting $\Delta\lambda_{LSPR}$ vs. analyte concentration, with concentration being plotted in the logarithmic scale in order to investigate non-specific adsorption at a lower concentration range.

The calibration curve equation was determined through linear regression on Origin software. Finally, the LOD was determined using a "Z value" of the blank, where Z = mean + 3σ, when σ = standard deviation of the blank. This Z value is obtained from six $\Delta\lambda_{LSPR}$ measurements using six different biosensors. The Z value is then inserted as the "Y" coordinate of the calibration curve equation, allowing the LOD concentration to be determined (Table S9). The concentrations of target microRNAs in patient samples were determined from the calibration curves developed in PBS buffer, with $\Delta\lambda_{LSPR}$ values and corresponding concentrations obtained from the average of six measurements. Each sample was independently analyzed in duplicate, and each sample UV-Visible spectra was the average of three individual scans conducted by the plate reader.

**BaseScope detection assay**

*Sample processing and pretreatment*

To define tissue expression patterns of miR-155, single molecule fluorescent in-situ hybridization (smFISH) was performed using a BaseScope duplex detection assay. Briefly, embedded pancreatic tissue sections from donors with T1D, AAb+ donors, and non-diabetic control donors were received from the Network of Pancreatic Organ Donors (nPOD) tissue bank (Table S10). The slides were baked at 60°C for 1 hour, then deparaffinized the tissue sections. This was accomplished by incubating the slides twice with xylene for 5 minutes and 100% ethanol for 2 minutes at room temperature (RT). The slides were air dried at 60°C for 5 minutes, then moved to RT, and treated with hydrogen peroxide (ACD Cell Diagnostics, CA, USA) for 10 minutes. After washing 5X with ddH2O, the slides were moved to a container with ddH2O at 99°C for 10 seconds, then immediately transferred to antigen-retrieval buffer (ACD Cell Diagnostics, CA, USA) at 99°C for 15 minutes. Next, slides were washed with ddH2O for 15 seconds at RT, followed by incubation with 100% ethanol for 3 minutes and air-drying at RT. Sections were applied with a hydrophobic barrier (Immedge, Vector labs) and air-dried at RT. The slides were transferred to a humidifying chamber and treated with protease IV (ACD Cell Diagnostics, CA, USA) for 30 minutes at 40°C in a HybEZ$^{TM}$ Oven (ACD Cell Diagnostics, CA, USA). Next, slides were washed 5X with ddH2O, and hybridization was performed according to the manufacturer's instructions using BaseScope duplex red assay probe sets.

*BaseScope assay and immunofluorescence staining*

The BaseScope duplex assay was performed per the instructions provided with the BaseScope kit protocol (ACD Cell Diagnostics, CA, USA). Because mature miRNAs are only ~22 nucleotides in length, we targeted to identify pre-miRNA expression in human tissue sections. Custom designed probe sets (ACD Cell Diagnostics, CA, USA) were generated to cover the regions of the human pre-miRNA sequence with a 1zz probe set covering 2-46 bp of pre-miR-155 (Cat# 18361A). Two control probe sets, PPIB-1zzas (Cat# 18282A, positive control) and bacterial DapB-1zz (Cat# 18267A, negative control), were used as internal controls to determine assay specificity. BaseScope Duplex Fast Red-B was used to detect pre-miR-155.

After the hybridization process was completed, the slides were blocked with donkey serum in PBS for 30 minutes, followed by immunofluorescent staining for human insulin, as reported previously (11,12). Nuclei staining was performed using DAPI (ACD Cell Diagnostics, CA, USA), and the slides were mounted with a coverslip in Prolong Gold Antifade mounting medium (Invitrogen, USA).

*Imaging*

Imaging was performed using a Zeiss LSM800 confocal microscopy attached to an Airyscan detector (Carl Zeiss, Germany). All the images were acquired using a 60X oil objective with Z-stack, then the images were processed using an Airyscan processer to determine the individual foci of the pre-miRNAs. As described above, four to five islets were randomly selected from each slide for imaging and were used to quantify miRNA expression.

**Machine learning-based smFISH image analysis.**

*Single-molecule quantification of miRNA in individual β cells*

Spatial quantification of pre-miRNA expression in single β cells was composed of two steps: (i) smFISH image processing and (ii) machine learning-based classification. We followed the protocol described in our previous publication to process the smFISH images of pre-miRNA-155 in human pancreatic islets (12). In brief, we segmented single nuclei in the DAPI channel via a binary Watershed algorithm in ImageJ (13). Next, we identified the cell-cell boundaries and the cytoplasm of single cells by dilating the nuclear mask with 150 pixels (5 um) in CellProfiler (14). The identification of β cells was accomplished by quantifying the fluorescent intensity in the corresponding insulin (Alexa 488) channel. Finally, the pre-miRNA was detected by a set of single particle detection algorithms, and the copy number was determined by normalizing the smFISH foci intensity ($I_{foci}$) by the intensity value of a single mRNA ($I_{mRNA}$). , ($N = I_{foci}/I_{mRNA}$). Each pre-miRNA localization was assigned to the nucleus or the cytoplasm of a given cell by encoding masked regions with a binary label and assigning the appropriate binary label based on transcript coordinates.

*Extracting the feature of miRNA distributions in a single β cell*

A single-cell phenotyping algorithm with supervised machine learning analyzed the spatial organization of the pre-miRNA in single β cells. Based on the preliminary processing and analysis of the pre-miRNA molecules, we utilized a series of descriptive methods, mathematical approaches, and statistical models to turn the spatial distribution of the pre-miRNA in a cell into an array of numbers. We then established a multi-dimensional feature library of the pre-miRNA in each β cell. This feature library consists of two categories of pre-miRNA features in a single cell.

The first group of features are descriptive quantifications of individual pre-miRNAs in a single cell, which include the distance of each pre-miRNA to the cell/nucleus centroid, the closest distance to the cell membrane/nucleic envelope, the distance of the pre-miRNA to its nearest neighbors, and other RNA localization-related properties (19). These values were averaged over all pre-miRNAs in a single cell, and each cell's mean value and standard deviation were kept as feature values. Meanwhile, we also included the spatial expression of the pre-miRNA as feature, which includes the number of pre-miRNAs in the cytoplasm or nucleus.

The two other groups in the feature library are the collective quantification of pre-miRNA location in a single cell. Two categories of quantification were used. First, Ripley's K function was used to describe the clustering effect of the pre-miRNA in a single cell (20). Ripley's K function quantifies the degree of clustering at a specific radius r by selecting individual RNA particles, then counting the average number of surrounding RNA particles that lie within the circle with a radius r. We then normalized and shifted Ripley's K function into Ripley's H function, which transforms the expectation value of a completely random distribution to 0. Therefore, a positive result of Ripley's H function at a specific radius r indicates a clustered distribution at the corresponding radius, and a negative result shows a dispersed distribution. Features are extracted by selecting Ripley's H function with specific intervals of radii. The extrema of Ripley's H function are also recorded as features . Secondly, the number of pre-miRNAs located at the nucleus and cell periphery was registered as boundary clustering features. This includes: i) nuclear pre-miRNA close to the nucleus envelope, ii) cytoplasmic pre-miRNA around the nucleus envelope, and iii) cytoplasmic pre-miRNA near the cellular membrane.

*Training Network and Classification.*

Once the feature library has been extracted from the cells, the features can then be used for classification: namely, the classification between healthy cells and cells from organ donors with AAb+ or established T1D. This classification analysis can determine if properties extracted from RNA localization serve as indicators for T1D or AAb+.

We used random forest classifier as the feature-based classification method (22,23), which notably can determine the importance of each feature, thus allowing us to identify the RNA localization properties with the strongest relationship with the specific cell conditions. The imbalanced-learn python library (24) was implemented to address the uneven number of cells in each sample by constructing each tree of the random forest classifier with a balanced bootstrap sample.

#### SUPPLEMENTAL FIGURES

Figure S1. Principal Component analysis on islets and EVs

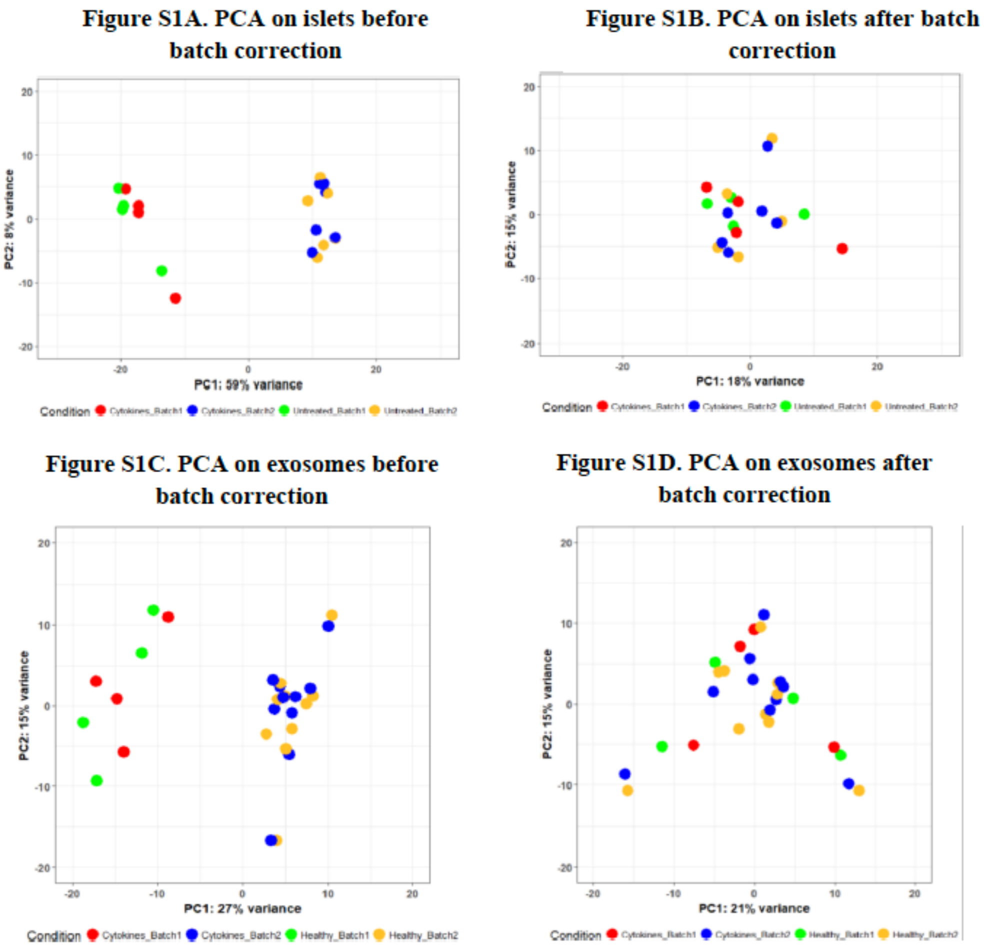

**Figure S1.** In both islets (S1A) and exosomes (S1C), principal component analysis using variance stabilized counts showed presence of batch effects as the samples clustered based on batches. PCA clustering using batch corrected variance stabilized counts revealed no clustering of samples based on batches, confirming removal of batch effects from both islets (S1B) and exosomes (S1D).

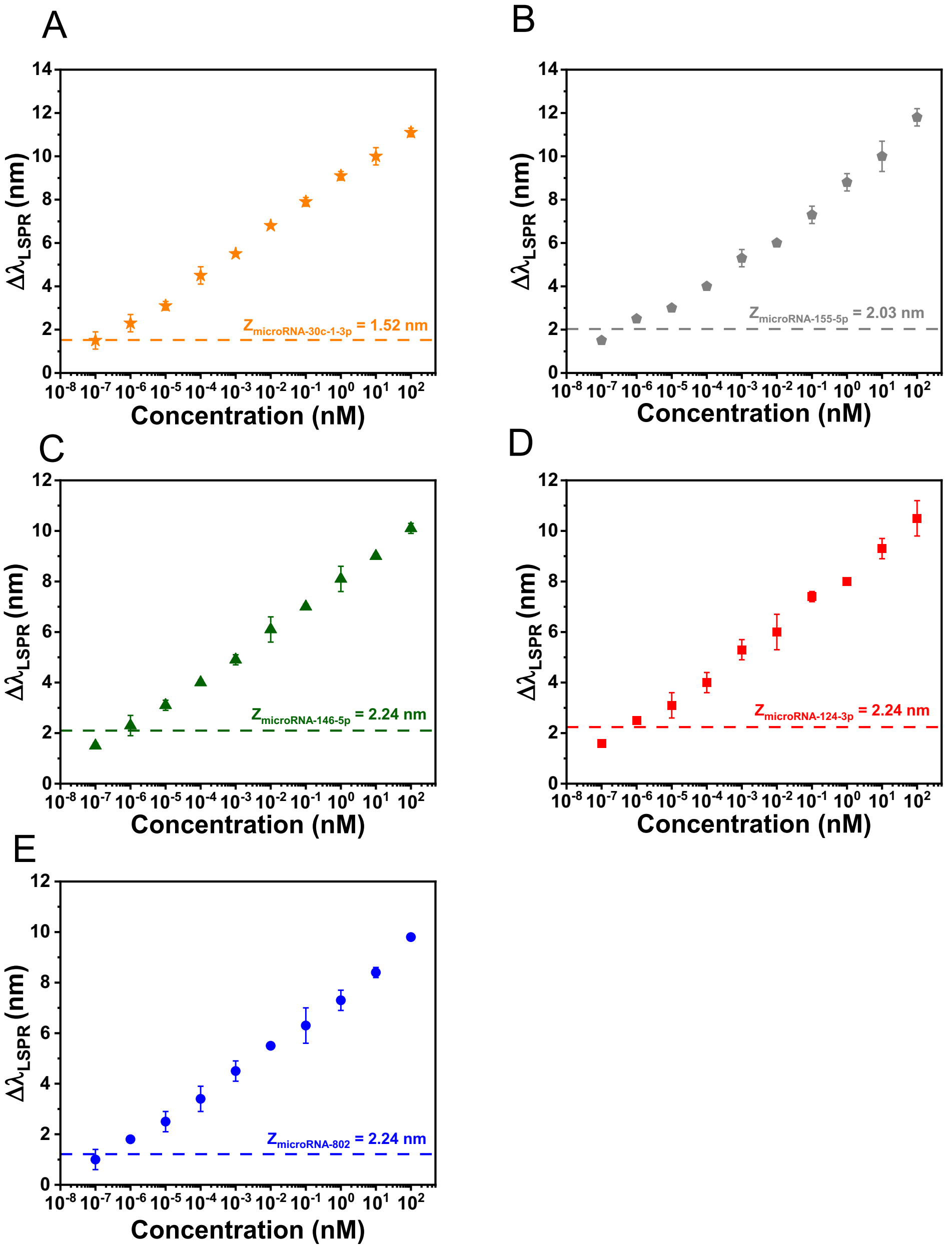

***Figure S2.*** *Calibration Curve of microRNAs (miR) representing $\Delta\lambda_{LSPR}$ vs.* Concentration (nM) being plotted in the logarithm scale *as described above;* ***A.*** *miR-30c-1-3p (orange star);* ***B.*** *mir-155-5p (gray pentagon);* ***C.*** *miR-146a-5p (green triangle);* ***D.*** *miR-124-3p (red square); and* ***E.*** *miR-802 (blue circle).*

**Figure S3.** Digital droplet PCR based determination of plasma derived EV-miRNAs

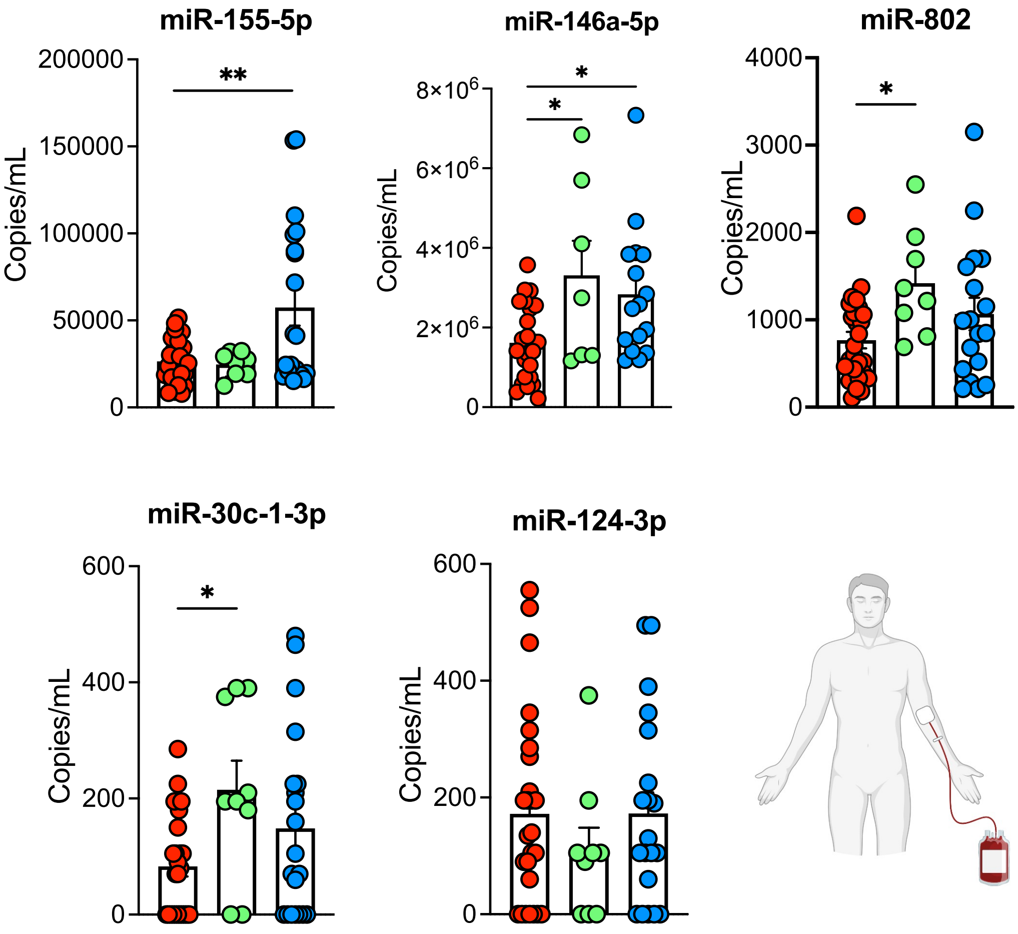

***Figure S3. ddPCR based measurement of plasma derived EV-miRNAs****. Plasma EV-miRNAs have been isolated from healthy controls (red), AAB+ (green) and new onset T1D (blue) and EV-miRNAs were measured using ddPCR and the data was represented as copies/mL.*

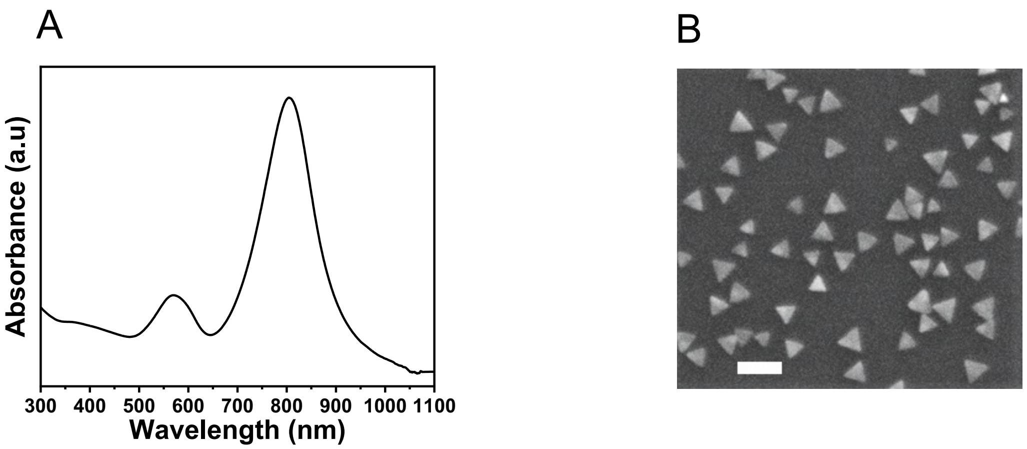

***Figure S4. (A)*** *UV-Visible absorption spectra of Au TNP solution in ACN (black, 801.0 nm);* ***(B)*** *A corresponding representative scanning electron microscope image of Au TNPs with a length of 42 nm and thickness of 8 nm. The scale bar is 100 nm. The dimension of the Au TNP was determined using ImageJ software.*

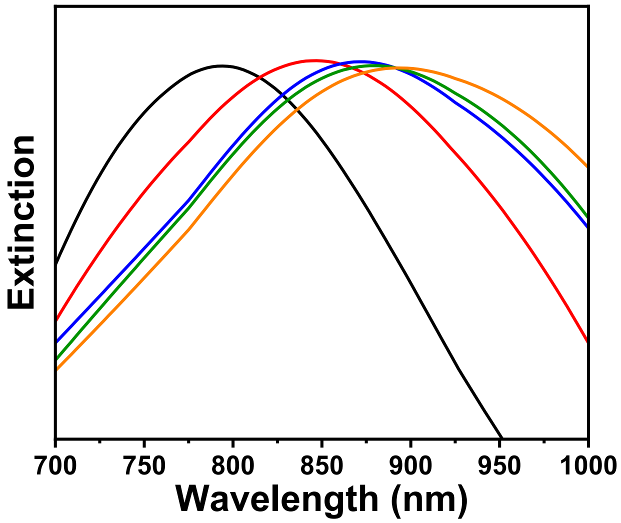

***Figure S5.*** *Representative UV-Vis extinction spectra showing Au TNPs before (black, $\lambda_{LSPR}$ = 799.9 nm), after functionalization with 75%:25% SP-HT:HT SAMs (red, $\lambda_{LSPR}$ = 853.0 nm), after exposure to UV light (blue, $\lambda_{LSPR}$ = 874.0 nm), followed by incubation in 10 µM - ssDNA-155-5p receptor in PBS buffer solution (green, $\lambda_{LSPR}$ = 886.2 nm) and lastly, incubation in 100 nM microRNA-155-5p in 10% human plasma (orange, 898.0 nm) .*

### SUPPLEMENTAL TABLES

#### Table S1

**Checklist for Reporting Human Islet Preparations Used in Research**

Adapted from Hart NJ, Powers AC (2018) Progress, challenges, and suggestions for using human islets to understand islet biology and human diabetes. Diabetologia https://doi.org/10.1007/s00125-018-4772-2.

Selected Islet Preparations for Carmella Evans Molina

Report Downloaded on 5/29/2023 3:48:39 PM

Data provided by IIDP

Labels in blue text are additional fields or clarification provided by IIDP

**Manuscript DOI:**

**Title: β Cell microRNAs Function as Molecular Hubs of Type 1 Diabetes Pathogenesis and as Biomarkers of Diabetes Risk**

**Author list: Farooq Syed, Preethi Krishnan, Garrick Chang, Sarah R. Langlais, Sumon Hati, Kentaro Yamada, Anh K. Lam, Sayali Talware, Xiaowen Liu, Rajesh Sardar, Jing Liu, Raghavendra G. Mirmira, and Carmella Evans-Molina**

**Corresponding author: Carmella Evans-Molina**

| Islet Preparation | 1 | 2 | 3 | 4 | 5 | 6 | 7 | 8 |
|---|---|---|---|---|---|---|---|---|
| **MANDATORY INFORMATION** | | | | | | | | |
| Unique identifier | SAMN08769073 | SAMN08769121 | SAMN08769129 | SAMN08769188 | SAMN08769191 | SAMN08769390 | SAMN09666985 | SAMN09704507 |
| Donor Age (years) | 28 | 28 | 44 | 30 | 35 | 33 | 48 | 61 |
| Donor Sex (M/F) | M | M | F | M | M | F | M | F |
| Donor BMI (kg/m$^2$) | 30.7 | 24.6 | 34.5 | 35.9 | 24.3 | 34.2 | 30.3 | 28.9 |
| Donor HbA1c | 4.9 | 5.1 | Not Documented | 5.4 | 5.4 | 5.5 | 5.8 | 5.2 |
| Origin/source of islets | IIDP | IIDP | IIDP | IIDP | IIDP | IIDP | IIDP | IIDP |
| Islet isolation center[a] | The Scharp-Lacy Research Institute | University of Pennsylvania | University of Pennsylvania | The Scharp-Lacy Research Institute | The Scharp-Lacy Research Institute | University of Wisconsin | The Scharp-Lacy Research Institute | The Scharp-Lacy Research Institute |
| Donor history of diabetes? Yes/No | No | No | No | No | No | No | No | No |
| **If Yes, complete the next two lines if this information is available** | | | | | | | | |
| Diabetes duration (years) | Not Applicable | Not Applicable | Not Applicable | Not Applicable | Not Applicable | Not Applicable | Not Applicable | Not Applicable |
| Glucose-lowering therapy at time of dea | Not Applicable | Not Applicable | Not Applicable | Not Applicable | Not Applicable | Not Applicable | Not Applicable | Not Applicable |
| **RECOMMENDED INFORMATION** | | | | | | | | |
| Cause of death | Cerebrovascular/stroke | Head trauma | Cerebrovascular/stroke | Cerebrovascular/stroke | Head trauma | Cerebrovascular/stroke | Anoxia | Head trauma |
| Warm ischemia time (h) | 0.1 | Not Reported | Not Reported | Not Reported | Not Reported | Not Reported | Not Reported | Not Reported |
| Cold ischemia time (h) | 7.4 | 13.4 | 8.4 | 11.6 | 7.4 | 14.3 | 11.2 | 6.2 |
| Estimated purity (%) By islet isolation center at time of shipment with Dithizone (DTZ) staining per IIDP SOP | 95 | 95 | 90 | 90 | 95 | 95 | 85 | 85 |
| Estimated viability (%) By islet isolation center at time of broadcast with Fluorescence Diacetate/Propidium Iodide (FDA/PI) staining per IIDP SOP | 95 | 91 | 90 | 95 | 95 | 98 | 95 | 95 |
| Total culture time(h)[c] Calculate[start date/time of experiment]-[date/ time islet culture began (below)] | To be completed by investigator | To be completed by investigator | To be completed by investigator | To be completed by investigator | To be completed by investigator | To be completed by investigator | To be completed by investigator | To be completed by investigator |
| Date/Time Islet Culture Began at islet isolation center (Pacific Standard Time, 24-hour format) | 2016-11-03 17:15 | 2016-10-06 15:00 | 2016-09-20 10:00 | 2016-07-24 18:35 | 2016-07-15 16:45 | 2016-05-29 18:22 | 2018-07-15 11:47 | 2018-07-19 15:33 |
| Glucose-stimulated insulin secretion or other functional measurement[d] Stimulation Index (SI) by static incubation at islet isolation center pre-shipment per IIDP SOP | SI (G2.8mM-G28mM)= 4.0 | SI (G2.8mM-G28mM)= 3.1 | SI (G2.8mM-G28mM)= 1.7 | SI (G2.8mM-G28mM)= 6.3 | SI (G2.8mM-G28mM)= 3.5 | SI (G2.8mM-G28mM)= 1.6 | SI (G2.8mM-G28mM)= 2.6 | SI (G2.8mM-G28mM)= 4.0 |
| Glucose-stimulated insulin secretion or other functional measurement Area Under the Curve (AUC) by perifusion at HIPP post shipment sample per HIPP SOP | AUC (G5.6mM-G16.7mM)= 40.4 (ng/100 IEQs) | AUC (G5.6mM-G16.7mM)= Not Available | AUC (G5.6mM-G16.7mM)= Not Availab | AUC (G5.6mM-G16.7mM)= Not Availab | AUC (G5.6mM-G16.7mM)= Not Availab | AUC (G5.6mM-G16.7mM)= Not Availab | AUC (G5.6mM-G16.7mM)= 46.3 (ng/10 | AUC (G5.6mM-G16.7mM)= 47.0 (ng/10 |
| Handpicked to purity? Yes/No | To be completed by investigator | To be completed by investigator | To be completed by investigator | To be completed by investigator | To be completed by investigator | To be completed by investigator | To be completed by investigator | To be completed by investigator |
| Additional notes | To be completed by investigator | To be completed by investigator | To be completed by investigator | To be completed by investigator | To be completed by investigator | To be completed by investigator | To be completed by investigator | To be completed by investigator |

| Islet Preparation | 9 | 10 | 11 | 12 | 13 | 14 | 15 | 16 |
|---|---|---|---|---|---|---|---|---|
| **MANDATORY INFORMATION** | | | | | | | | |
| Unique identifier | SAMN09767131 | SAMN09767462 | SAMN10252228 | SAMN11155033 | SAMN16871526 | SAMN17098146 | SAMN17367763 | SAMN28867622 |
| Donor Age (years) | 29 | 44 | 55 | 51 | 62 | 57 | 56 | 36 |
| Donor Sex (M/F) | F | M | F | M | M | M | M | M |
| Donor BMI (kg/m$^2$) | 24.1 | 29.8 | 35.7 | 24.0 | 21.2 | 24.8 | 33.0 | 29.6 |
| Donor HbA1c | 4.3 | 5.4 | 6.0 | 4.7 | 5.3 | 5.9 | 5.3 | 5.4 |
| Origin/source of islets | IIDP | IIDP | IIDP | IIDP | IIDP | IIDP | IIDP | IIDP |
| Islet isolation center[a] | Southern California Islet Cell Resource C | University of Wisconsin | University of Wisconsin | Southern California Islet Cell Resource C | The Scharp-Lacy Research Institute | Southern California Islet Cell Resource C | The Scharp-Lacy Research Institute | Southern California Islet Cell Resource Center |
| Donor history of diabetes? Yes/No | No | No | No | No | No | No | No | No |
| **If Yes, complete the next two lines if this information is available** | | | | | | | | |
| Diabetes duration (years) | Not Applicable | Not Applicable | Not Applicable | Not Applicable | Not Applicable | Not Applicable | Not Applicable | Not Applicable |
| Glucose-lowering therapy at time of dea | Not Applicable | Not Applicable | Not Applicable | Not Applicable | Not Applicable | Not Applicable | Not Applicable | Not Applicable |
| **RECOMMENDED INFORMATION** | | | | | | | | |
| Cause of death | Cerebrovascular/stroke | Head trauma | Cerebrovascular/stroke | Head trauma | Cerebrovascular/stroke | Cerebrovascular/stroke | Head trauma | Anoxia |
| Warm ischemia time (h) | Not Reported | Not Reported | Not Reported | Not Reported | Not Reported | 0.4 | 0.7 | 0.4 |
| Cold ischemia time (h) | 7.0 | 12.5 | 8.2 | 8.8 | 9.7 | 8.1 | 10.9 | 8.2 |
| Estimated purity (%) By islet isolation center at time of shipment with Dithizone (DTZ) staining per IIDP SOP | 80 | 90 | 93 | 80 | 90 | 85 | 90 | 93 |
| Estimated viability (%) By islet isolation center at time of broadcast with Fluorescence Diacetate/Propidium Iodide (FDA/PI) staining per IIDP SOP | 96 | 95 | 98 | 96 | 95 | 98 | 95 | 96 |
| Total culture time(h)[c] Calculate[start date/time of experiment]-[date/ time islet culture began (below)] | To be completed by investigator | To be completed by investigator | To be completed by investigator | To be completed by investigator | To be completed by investigator | To be completed by investigator | To be completed by investigator | To be completed by investigator |
| Date/Time Islet Culture Began at islet isolation center (Pacific Standard Time, 24-hour format) | 2018-08-03 16:40 | 2018-08-05 22:30 | 2018-10-16 19:30 | 2019-03-15 13:40 | 2020-11-20 04:00 | 2020-12-16 13:20 | 2021-01-15 14:30 | 2022-06-04 12:15 |
| Glucose-stimulated insulin secretion or other functional measurement[d] Stimulation Index (SI) by static incubation at islet isolation center pre-shipment per IIDP SOP | SI (G2.8mM-G28mM)= 2.8 | SI (G2.8mM-G28mM)= 1.7 | SI (G2.8mM-G28mM)= 3.8 | SI (G2.8mM-G28mM)= 1.0 | SI (G2.8mM-G28mM)= 1.1 | SI (G2.8mM-G28mM)= 1.2 | SI (G2.8mM-G28mM)= 1.5 | SI (G2.8mM-G28mM)= 1.8 |
| Glucose-stimulated insulin secretion or other functional measurement Area Under the Curve (AUC) by perifusion at HIPP post shipment sample per HIPP SOP | AUC (G5.6mM-G16.7mM)= 34.3 (ng/10 | AUC (G5.6mM-G16.7mM)= 16.4 (ng/10 | AUC (G5.6mM-G16.7mM)= 27.8 (ng/10 | AUC (G5.6mM-G16.7mM)= Not Availab | AUC (G5.6mM-G16.7mM)= 18.9 (ng/10 | AUC (G5.6mM-G16.7mM)= -10.0 (ng/1 | AUC (G5.6mM-G16.7mM)= 51.9 (ng/10 | AUC (G5.6mM-G16.7mM)= 18.5 (ng/100 IEQs) |
| Handpicked to purity? Yes/No | To be completed by investigator | To be completed by investigator | To be completed by investigator | To be completed by investigator | To be completed by investigator | To be completed by investigator | To be completed by investigator | To be completed by investigator |
| Additional notes | To be completed by investigator | To be completed by investigator | To be completed by investigator | To be completed by investigator | To be completed by investigator | To be completed by investigator | To be completed by investigator | To be completed by investigator |

#### Table S1 (Continued)

| 17 | 18 | 19 | 20 | 21 | 22 | 23 | 24 |
|---|---|---|---|---|---|---|---|
| SAMN08769121 | SAMN08769124 | SAMN08769132 | SAMN08769199 | SAMN08769206 | SAMN08769390 | SAMN08769836 | SAMN08773446 |
| 28 | 15 | 23 | 43 | 52 | 33 | 56 | 69 |
| M | M | M | F | F | F | F | F |
| 24.6 | 24.6 | 24.8 | 29.5 | 25.8 | 34.2 | 33.5 | 22.1 |
| 5.1 | 5.1 | 5.3 | 5.7 | 5.6 | 5.5 | 5.2 | 5.3 |
| IIDP | IIDP | IIDP | IIDP | IIDP | IIDP | IIDP | IIDP |
| University of Pennsylvania | Southern California Islet Cell Resource Center | The Scharp-Lacy Research Institute | Southern California Islet Cell Resource Center | University of Miami | University of Wisconsin | The Scharp-Lacy Research Institute | The Scharp-Lacy Research Institute |
| No | No | No | No | No | No | No | No |
| Not Applicable | Not Applicable | Not Applicable | Not Applicable | Not Applicable | Not Applicable | Not Applicable | Not Applicable |
| Not Applicable | Not Applicable | Not Applicable | Not Applicable | Not Applicable | Not Applicable | Not Applicable | Not Applicable |
| Head trauma | Head trauma | Anoxia | Cerebrovascular/stroke | Cerebrovascular/stroke | Cerebrovascular/stroke | HEAD TRAUMA | Cerebrovascular/stroke |
| Not Reported | Not Reported | Not Reported | 0.0 | Not Reported | Not Reported | Not Reported | Not Reported |
| 13.4 | Not Documented | 8.2 | Not Documented | 6.3 | 14.3 | 11.4 | 9.2 |
| 95 | 80 | 95 | 80 | 90 | 95 | 90 | 90 |
| 91 | 97 | 95 | 97 | 95 | 98 | 95 | 95 |
| To be completed by investigator | To be completed by investigator | To be completed by investigator | To be completed by investigator | To be completed by investigator | To be completed by investigator | To be completed by investigator | To be completed by investigator |
| 2016-10-06 15:00 | 2016-09-30 11:00 | 2016-09-08 18:34 | 2016-06-30 15:00 | 2016-08-11 06:50 | 2016-05-29 18:22 | 2016-01-15 22:15 | 2016-04-01 22:30 |
| SI (G2.8mM-G28mM)= 3.1 | SI (G2.8mM-G28mM)= 2.5 | SI (G2.8mM-G28mM)= 3.5 | SI (G2.8mM-G28mM)= 1.8 | SI (G2.8mM-G28mM)= 2.7 | SI (G2.8mM-G28mM)= 1.6 | SI (G2.8mM-G28mM)= 5.3 | SI (G2.8mM-G28mM)= 2.3 |
| AUC (G5.6mM-G16.7mM)= Not Available | AUC (G5.6mM-G16.7mM)= 35.1 (ng/100 IEQs) | AUC (G5.6mM-G16.7mM)= 24.8 (ng/100 IEQs) | AUC (G5.6mM-G16.7mM)= 41.1 (ng/100 IEQs) | AUC (G5.6mM-G16.7mM)= Not Available | AUC (G5.6mM-G16.7mM)= Not Available | AUC (G5.6mM-G16.7mM)= Not Available | AUC (G5.6mM-G16.7mM)= Not Available |
| To be completed by investigator | To be completed by investigator | To be completed by investigator | To be completed by investigator | To be completed by investigator | To be completed by investigator | To be completed by investigator | To be completed by investigator |
| To be completed by investigator | To be completed by investigator | To be completed by investigator | To be completed by investigator | To be completed by investigator | To be completed by investigator | To be completed by investigator | To be completed by investigator |

| 25 | 26 | 27 | 28 | 29 | 30 | 31 | 32 |
|---|---|---|---|---|---|---|---|
| SAMN08774476 | SAMN08774480 | SAMN08774814 | SAMN08768705 | SAMN08769076 | SAMN08769132 | SAMN19796386 | |
| 39 | 24 | 45 | 27 | 39 | 23 | 40 | 19 |
| F | F | F | F | M | M | M | M |
| 22.8 | 35.4 | 32.8 | 27.7 | 23.1 | 24.8 | 31.7 | 20.9 |
| 6.0 | Not Done | Not Done | 5.0 | 4.5 | 5.3 | | |
| IIDP | IIDP | IIDP | IIDP | IIDP | IIDP | ADI | NDRI |
| University of Miami | University of Miami | University of Wisconsin | The Scharp-Lacy Research Institute | The Scharp-Lacy Research Institute | The Scharp-Lacy Research Institute | Alberta Diabetes Institute | NDRI |
| No | No | No | No | No | No | No | |
| Not Applicable | Not Applicable | Not Applicable | Not Applicable | Not Applicable | Not Applicable | Not Applicable | Not Applicable |
| Not Applicable | Not Applicable | Not Applicable | Not Applicable | Not Applicable | Not Applicable | Not Applicable | Not Applicable |
| Cerebrovascular/stroke | Cerebrovascular/stroke | Cerebrovascular/stroke | Anoxia | Cerebrovascular/stroke | Anoxia | NDD-Neurological | Head trauma |
| Not Reported | Not Reported | Not Reported | Not Reported | Not Reported | Not Reported | Not Reported | |
| 23.4 | 12.8 | 3.8 | 14.4 | 6.2 | 8.2 | 4 | |
| 85 | 90 | 90 | 90 | 90 | 95 | 75 | 90 |
| 93 | 92 | 98 | 95 | 95 | 95 | | 95 |
| To be completed by investigator | To be completed by investigator | To be completed by investigator | To be completed by investigator | To be completed by investigator | To be completed by investigator | | |
| 2015-03-14 08:30 | 2015-02-27 13:00 | 2015-02-17 04:00 | 2017-01-12 15:35 | 2016-10-26 16:11 | 2016-09-08 18:34 | | |
| SI (G2.8mM-G28mM)= 5.1 | SI (G2.8mM-G28mM)= 3.7 | SI (G2.8mM-G28mM)= 2.3 | SI (G2.8mM-G28mM)= 2.5 | SI (G2.8mM-G28mM)= 5.7 | SI (G2.8mM-G28mM)= 3.5 | | |
| AUC (G5.6mM-G16.7mM)= Not Available | AUC (G5.6mM-G16.7mM)= Not Available | AUC (G5.6mM-G16.7mM)= Not Available | AUC (G5.6mM-G16.7mM)= 36.4 (ng/100 IEQs) | AUC (G5.6mM-G16.7mM)= 55.7 (ng/100 IEQs) | AUC (G5.6mM-G16.7mM)= 24.8 (ng/100 IEQs) | | |
| To be completed by investigator | To be completed by investigator | To be completed by investigator | To be completed by investigator | To be completed by investigator | To be completed by investigator | | |
| To be completed by investigator | To be completed by investigator | To be completed by investigator | To be completed by investigator | To be completed by investigator | To be completed by investigator | | |

**Table S2. Descriptive statistics**

**Table S2A. Descriptive statistics of islets**

| Sample name | Batch | Treatment | Total reads | Total aligned reads | Total unaligned reads | Mapped to miRNA |
|---|---|---|---|---|---|---|
| Sample 1_Cytokine | 1 | Cytokine | 2,054,411 | 1,300,648 | 753,764 | 660,907 |
| Sample 2_Cytokine | 1 | Cytokine | 3,124,061 | 2,127,798 | 996,377 | 1,091,088 |
| Sample 3_Cytokine | 1 | Cytokine | 2,591,878 | 1,694,829 | 897,065 | 656,258 |
| Sample 4_Cytokine | 1 | Cytokine | 2,326,519 | 1,664,624 | 661,908 | 622,900 |
| Sample 5_Cytokine | 2 | Cytokine | 14,664,373 | 12,065,846 | 2,599,253 | 2,451,914 |
| Sample 6_Cytokine | 2 | Cytokine | 13,377,553 | 9,496,725 | 3,880,288 | 2,272,287 |
| Sample 7_Cytokine | 2 | Cytokine | 8,823,179 | 5,180,088 | 3,642,957 | 1,571,907 |
| Sample 8_Cytokine | 2 | Cytokine | 14,697,258 | 10,689,316 | 4,007,947 | 2,038,327 |
| Sample 9_Cytokine | 2 | Cytokine | 15,187,948 | 10,303,504 | 4,883,966 | 1,853,587 |
| Sample 10_Cytokine | 2 | Cytokine | 11,132,752 | 6,025,045 | 5,107,693 | 1,754,292 |
| Sample 1_untreated | 1 | Untreated | 1,854,137 | 945,981 | 908,195 | 498,742 |
| Sample 2_untreated | 1 | Untreated | 2,352,048 | 1,179,317 | 1,172,746 | 582,585 |
| Sample 3_untreated | 1 | Untreated | 2,401,487 | 1,357,801 | 1,043,636 | 553,268 |
| Sample 4_untreated | 1 | Untreated | 2,349,003 | 1,757,994 | 591,095 | 928,985 |
| Sample 5_untreated | 2 | Untreated | 13,132,128 | 9,988,297 | 3,144,136 | 2,616,396 |
| Sample 6_untreated | 2 | Untreated | 14,963,188 | 11,017,395 | 3,946,395 | 1,871,802 |
| Sample 7_untreated | 2 | Untreated | 16,269,364 | 11,944,967 | 4,325,159 | 1,889,477 |
| Sample 8_untreated | 2 | Untreated | 12,869,993 | 8,602,303 | 4,267,563 | 2,316,388 |
| Sample 9_untreated | 2 | Untreated | 6,662,954 | 4,308,266 | 2,354,925 | 839,713 |

| | | | | | | |
|---|---|---|---|---|---|---|
| Sample 10_untreated | 2 | Untreated | 10,287,444 | 6,178,639 | 4,108,689 | 1,332,384 |

**Table S2B. Descriptive statistics of EVs**

| Sample name | Batch | Treatment | Total reads | Total aligned | Total unaligned | Mapped to miRNAs |
|---|---|---|---|---|---|---|
| Sample 1_Cytokine | 2 | Cytokine | 4,264,849 | 2,863,420 | 1,401,544 | 1,176,172 |
| Sample 2_Cytokine | 2 | Cytokine | 2,135,118 | 1,082,718 | 1,052,459 | 255,727 |
| Sample 3_Cytokine | 2 | Cytokine | 4,055,979 | 2,141,963 | 1,913,961 | 557,978 |
| Sample 4_Cytokine | 2 | Cytokine | 3,356,133 | 2,013,344 | 1,342,806 | 569,917 |
| Sample 5_Cytokine | 2 | Cytokine | 3,697,693 | 1,974,198 | 1,723,462 | 510,295 |
| Sample 6_Cytokine | 1 | Cytokine | 3,373,475 | 1,981,579 | 1,391,753 | 499,652 |
| Sample 7_Cytokine | 2 | Cytokine | 4,372,133 | 1,929,860 | 2,442,437 | 372,539 |
| Sample 8_Cytokine | 1 | Cytokine | 2,199,842 | 1,413,398 | 786,369 | 393,698 |
| Sample 9_Cytokine | 2 | Cytokine | 5,132,436 | 3,181,084 | 1,951,577 | 1,045,087 |
| Sample 10_Cytokine | 1 | Cytokine | 1,031,149 | 709,327 | 321,805 | 167,226 |
| Sample 11_Cytokine | 2 | Cytokine | 3,757,155 | 2,037,129 | 1,719,916 | 786,489 |
| Sample 12_Cytokine | 1 | Cytokine | 3,919,830 | 2,854,420 | 1,065,575 | 806,124 |
| Sample 13_Cytokine | 2 | Cytokine | 4,000,971 | 2,270,551 | 1,730,429 | 1,084,639 |
| Sample 14_Cytokine | 2 | Cytokine | 3,606,848 | 1,511,991 | 2,094,851 | 190,957 |
| Sample 1_untreated | 2 | Healthy | 4,095,296 | 3,139,454 | 955,906 | 1,493,497 |
| Sample 2_untreated | 2 | Healthy | 2,222,263 | 1,080,686 | 1,141,621 | 227,017 |
| Sample 3_untreated | 2 | Healthy | 3,136,268 | 1,767,914 | 1,368,472 | 414,766 |
| Sample 4_untreated | 2 | Healthy | 2,200,147 | 1,184,119 | 1,016,069 | 425,164 |
| Sample 5_untreated | 2 | Healthy | 2,687,655 | 1,265,886 | 1,421,779 | 275,131 |
| Sample 6_untreated | 1 | Healthy | 2,482,904 | 1,052,503 | 1,430,455 | 211,606 |
| Sample 7_untreated | 2 | Healthy | 2,726,129 | 1,270,376 | 1,455,826 | 228,409 |
| Sample 8_untreated | 1 | Healthy | 2,013,477 | 1,265,470 | 747,934 | 406,393 |
| Sample 9_untreated | 2 | Healthy | 3,139,816 | 1,711,828 | 1,428,073 | 669,261 |
| Sample 10_untreated | 1 | Healthy | 1,028,515 | 628,423 | 400,084 | 98,767 |
| Sample 11_untreated | 2 | Healthy | 3,744,869 | 1,717,022 | 2,027,995 | 424,485 |
| Sample 12_untreated | 1 | Healthy | 4,428,561 | 2,899,822 | 1,528,622 | 981,547 |
| Sample 13_untreated | 2 | Healthy | 3,525,663 | 1,623,568 | 1,902,017 | 491,683 |
| Sample 14_untreated | 2 | Healthy | 3,769,590 | 1,778,493 | 1,990,927 | 240,566 |

**Table S3. Differentially expressed miRNAs in islets and EVs.** a fold change of ≥ 1.5 and p < 0.05 were considered as differentially expressed. 20 miRNAs and 14 miRNAs were identified as differentially expressed from islets (left panel) and EVs (right panel), respectively. miRNAs highlighted in red indicate miRNAs common between islets and extracellular vesicles.

| Differentially expressed miRNAs in islets | | | Differentially expressed miRNAs in EVs | | |
|---|---|---|---|---|---|
| **miRNA** | **Linear scale fold change** | **p-value** | **miRNA** | **Linear scale fold change** | **p-value** |
| hsa-miR-155-5p | 7.13 | 4.07E-40 | hsa-miR-155-5p | 2.59 | 0.0001 |
| hsa-miR-6891-5p | 4.58 | 2.41E-08 | hsa-miR-1226-3p | 2.02 | 0.0178 |
| hsa-miR-4640-5p | 2.47 | 0.0002 | hsa-miR-802 | 1.63 | 1.64E-05 |
| hsa-miR-146a-5p | 2.4 | 1.15E-09 | hsa-miR-146a-5p | 1.63 | 2.97E-05 |
| hsa-miR-205-5p | 2.29 | 1.13E-07 | hsa-miR-31-3p | 1.6 | 0.0085 |
| hsa-miR-338-3p | 2.15 | 0.0113 | hsa-miR-30c-1-3p | 1.6 | 0.0480 |
| hsa-miR-21-3p | 2.08 | 3.59E-08 | hsa-miR-193b-5p | 1.56 | 0.0031 |
| hsa-miR-27a-5p | 2.03 | 0.0052 | hsa-miR-193b-3p | 1.53 | 0.0004 |
| hsa-miR-6743-3p | 1.68 | 0.0479 | hsa-miR-148a-3p | 1.52 | 4.30E-05 |
| hsa-miR-23a-5p | 1.63 | 0.0008 | hsa-miR-217 | 1.51 | 0.0001 |
| hsa-miR-431-5p | 1.61 | 0.0006 | hsa-miR-216a-3p | 1.51 | 0.0085 |
| hsa-miR-363-3p | 1.57 | 0.0260 | hsa-miR-4485 | -1.67 | 0.0111 |
| hsa-miR-194-3p | 1.52 | 0.0047 | hsa-miR-210-5p | -1.8 | 0.0436 |
| hsa-miR-147b | 1.52 | 0.0080 | hsa-miR-124-3p | -2.22 | 0.0092 |
| hsa-miR-543 | 1.5 | 0.0001 | | | |
| hsa-miR-33a-5p | -1.55 | 0.0113 | | | |
| hsa-miR-181a-2-3p | -1.8 | 3.14E-05 | | | |
| hsa-miR-335-3p | -1.88 | 0.0094 | | | |
| hsa-miR-675-5p | -2.65 | 5.36E-06 | | | |
| hsa-miR-338-5p | -3.12 | 0.0042 | | | |

### Table S4: ddPCR control gene synthesis

| Name | ID | Matured miRNA | Length (bp) | DNA sequence converted from miRNA (U --> T) | Matured miRNA + flanking random DNA sequence, 100 bp on both sides, totally ~220-224 bp |
|---|---|---|---|---|---|
| | hsa-miR-155-5p MIMAT0000646 | UUAAUGCUAAUCGUGAUAGGGGUU | 24 | TTAATGCTAATCGTGATAGGGGTT | ctgacatcgcggggtcatgcactagtggtaccgatggcgtcttgagacttctaggatctaattggtgccccctcccgttttaaatccatcgttactccccTTAATGCTAATCGTGATAGGGGTTtccgccttttaatagagcttataggaaagaatacagtacgtacgggtgtggagactagtggagcttaactctgatagcgtgtgaggtaaggttagagcag |
| | hsa-miR-146a-5p MIMAT0000449 | UGAGAACUGAAUUCCAUGGGUU | 22 | TGAGAACTGAATTCCATGGGTT | gggtcgtatacggcatcggatctacgctgagattaggcattaataccagattagcaattaaaatctcatagctcattgggatttctcttcttaacaactaTGAGAACTGAATTCCATGGGTTaatggtttagctcacccttagtacctcctcgatgcgtttatgcaccacgtggcgtggggggttacccgtctacatttacggcgctcctgcagataccaat |
| | hsa-miR-802 MIMAT0004185 | CAGUAACAAAGAUUCAUCCUUGU | 23 | CAGTAACAAAGATTCATCCTTGT | ctggcgcccataggcaaacttacacaatcgacaagcggattcatgatgcagcaagcctaggcatcatttctctatcgtgtgggactcgatcttcgccgcaCAGTAACAAAGATTCATCCTTGTacattctggaagaccatatctcatggctggcccctttgagcgctgaaatcaaagagcacgatggttatccatatagcttgcgcacgtaattctcgttgt |
| | hsa-miR-124-3p MIMAT0000422 | UAAGGCACGCGGUGAAUGCCAA | 22 | TAAGGCACGCGGTGAATGCCAA | ttaataaaagctgaatgaacatctcgaggtgcccagcggatggaaactatgcctctgcttaacccgtgtttcttccccgactgtacacatgtgtagttctTAAGGCACGCGGTGAATGCCAAggcatggtaactgaccgctcttgatgcgatacgagggcacatgacgttcccttgagagaactagacggagaaacccgcaccgtgaaatcagttggctata |
| | hsa-miR-30c-1-3p MIMAT0004674 | CUGGGAGAGGGUUGUUUACUCC | 22 | CTGGGAGAGGGTTGTTTACTCC | taatcaacgacactaaaaaacagacttcatgccgaaaccttgtgcagttgcttcgggcgcgcacccaatgatgcacgtattacctgtcacaagtcggaatCTGGGAGAGGGTTGTTTACTCCtcgttgtcgaagaatacttgtgcttattaaaggaaggttccagccggcctacaaacgttttttggtttaatgtgtgcgttggggtccggtatatggcgca |
| | mmu-miR-155-5p MIMAT0000165 | UUAAUGCUAAUUGUGAUAGGGGU | 23 | TTAATGCTAATTGTGATAGGGGT | gctcagctcagatccccatttcctcctcgggatacacgatcgtgcccctgagatctagactatcgctcccctctctcgcgattcggcccgccggcgcttgTTAATGCTAATTGTGATAGGGGTtttgatgctttctactcttcgcgtaatatggactcaaggccgaatcgttatgctgcccgagtggcaatcggtcttcgggcggaattcgcatacaagagac |
| | mmu-miR-124-3p MIMAT0000134 | UAAGGCACGCGGUGAAUGCC | 20 | TAAGGCACGCGGTGAATGCC | gtctcaatgcctggtaaaatggctgaatcagatacgagtcgcgaatgcttagaatgagttgtgtaacttagagtaccccagcgcagatgtatgtaccgttTAAGGCACGCGGTGAATGCCgtatacaggcctggcaaaggtgctcgtggtctcgtcgcaggtttcgcagtctaaggagctcttcgaccgaattacgcgtctccctctgacccacagcatg |
| | mmu-miR-802-5p MIMAT0004188 | UCAGUAACAAAGAUUCAUCCUU | 22 | TCAGTAACAAAGATTCATCCTT | tccgagtaacctgatctaaaccgggacagcttactagcgaggtataacgatccgcagcgacaatgatctaagccacgcaccattaaaatcgcctcatatgTCAGTAACAAAGATTCATCCTTactacatcctaagatgatgtgctagagggatgcgtccctatcgtctaaggctggatggacctacggtggcccattttaacaatacgcatagcttgcacac |

**Procedures**

1 miRNA sequence was converted to DNA sequence using the program on the website (http://biomodel.uah.es/en/lab/cybertory/analysis/trans.htm)

2 Random DNA sequences were generated using the program on the GenScript website (https://www.genscript.com/sms2/random_dna.html)

3 Put 100 bp random DNA sequences onto both ends of miRNA sequences

4 Run NCBI-BLAST and check whether there is any overlap between the designed sequences and any known sequences except the target miRNA sequences

**Table S5. Single stranded DNA (*-ssDNA)* oligomer sequences used in this study.**

| Name | Nucleic Acid -ssDNA Sequence |
|---|---|
| -ssDNA-155-5p | 5' - AAC CCC TAT CAC GAT TAG CAT TAA – 3' |
| -ssDNA-146a-5p | 5' – AAC CCA TGG AAT TCA GTT CTC A – 3' |
| -ssDNA-802 | 5' - ACA AGG ATG AAT CTT TGT TAC TG – 3' |
| -ssDNA-124-3p | 5' - TTG GCA TTC ACC GCG TGC CTT A – 3' |
| -ssDNA-30c-1-3p | 5' - GGA GTA AAC AAC CCT CTC CCA G – 3' |

**Table S6.** microRNA sequences that were used to generate calibration curves.

| Name | Nucleic Acid microRNA Sequence |
|---|---|
| miR-155-5p | 5’- UUA AUG CUA AUC GUG AUA GGG GUU -3’ |
| miR-146a-5p | 5’ – UGA GAA CUG AAU UCC AUG GGU U – 3’ |
| miR-802 | 5’ - CAG UAA CAA AGA UUC AUC CUU GU – 3’ |
| miR-124-3p | 5’ - UAA GGC ACG CGG UGA AUG CCA A – 3’ |
| miR-30c-1-3p | 5’ – CUG GGA GAG GGU UGU UUA CUC C – 3’ |

**Table S7.** $\Delta\lambda_{LSPR}$ responses of spiropyran based plasmonic nanosensors for microRNA-155-5p, -802, -124-3p, 30c-1-3p and -146a-5p in human plasma.

| microRNA | microRNA Concentration (nM) | Average $\Delta\lambda_{LSPR}$ (nm) | Standard Deviation (nm) |
|---|---|---|---|
| miR-155-5p | 100 | 11.8 | 0.4 |
| | 10 | 10.0 | 0.7 |
| | 1 | 8.8 | 0.4 |
| | 0.1 | 7.3 | 0.4 |
| | 0.01 | 6.0 | 0.1 |
| | 0.001 | 5.3 | 0.4 |
| | 0.0001 | 4.0 | 0.1 |
| | 0.00001 | 3.0 | 0.1 |
| | 0.000001 | 2.5 | 0.1 |
| | 0.0000001 | 1.5 | 0.1 |
| miR-124-3p | 100 | 10.5 | 0.7 |
| | 10 | 9.3 | 0.4 |
| | 1 | 8.0 | 0.1 |
| | 0.1 | 7.4 | 0.2 |
| | 0.01 | 6.0 | 0.7 |
| | 0.001 | 5.3 | 0.4 |
| | 0.0001 | 4.0 | 0.4 |
| | 0.00001 | 3.1 | 0.5 |
| | 0.000001 | 2.5 | 0.1 |
| | 0.0000001 | 1.6 | 0.1 |
| miR-802 | 100 | 9.8 | 0.1 |
| | 10 | 8.4 | 0.2 |
| | 1 | 7.3 | 0.4 |
| | 0.1 | 6.3 | 0.7 |
| | 0.01 | 5.5 | 0.1 |
| | 0.001 | 4.5 | 0.4 |
| | 0.0001 | 3.4 | 0.5 |
| | 0.00001 | 2.5 | 0.4 |
| | 0.000001 | 1.8 | 0.1 |
| | 0.0000001 | 1.0 | 0.4 |
| miR-146a-5p | 100 | 10.1 | 0.2 |
| | 10 | 9.0 | 0.1 |
| | 1 | 8.1 | 0.5 |
| | 0.1 | 7.0 | 0.1 |
| | 0.01 | 6.1 | 0.5 |
| | 0.001 | 4.9 | 0.2 |
| | 0.0001 | 4.0 | 0.1 |
| | 0.00001 | 3.1 | 0.2 |
| | 0.000001 | 2.3 | 0.4 |
| | 0.0000001 | 1.5 | 0.1 |
| miR-30c-1-3p | 100 | 11.1 | 0.2 |
| | 10 | 10.0 | 0.4 |
| | 1 | 9.1 | 0.2 |
| | 0.1 | 7.9 | 0.2 |
| | 0.01 | 6.8 | 0.1 |
| | 0.001 | 5.5 | 0.1 |
| | 0.0001 | 4.5 | 0.4 |
| | 0.00001 | 3.1 | 0.2 |
| | 0.000001 | 2.3 | 0.4 |
| | 0.0000001 | 1.5 | 0.4 |

**Table S8.** $\Delta\lambda_{LSPR}$ responses of spiropyran based plasmonic nanosensors for nucleic acid -ssDNA attachment and microRNA targets at 100 nM concentrations.

| Nucleic Acid Gene | Average $\Delta\lambda_{LSPR}$ (nm) after attachment of corresponding 10 µM -ssDNA receptor | Average $\Delta\lambda_{LSPR}$ (nm) after addition of 100 nM Nucleic Acid microRNA Target |
|---|---|---|
| **hsa-155-5p gene** | *+12.0 (±0.9)* | *+11.8 (±0.4)* |
| **hsa-146a-5p gene** | *+11.6 (±1.9)* | *+10.1 (±0.2)* |
| **hsa-802 Gene** | *+10.9 (±1.7)* | *+9.8 (±0.1)* |
| **hsa-124-3p Gene** | *+11.8 (±1.0)* | *+10.5 (±0.7)* |
| **hsa-30c-1-3p Gene** | *+12.2 (±1.6)* | *+11.1 (±0.2)* |
